# RTK signalling promotes epithelial columnar cell shape and apical junction maintenance in human lung progenitor cells

**DOI:** 10.1101/2022.09.08.507126

**Authors:** Shuyu Liu, Dawei Sun, Richard Butler, Emma L. Rawlins

**Affiliations:** Wellcome Trust/CRUK Gurdon Institute, University of Cambridge, Cambridge, CB2 1QN, UK.; Department of Physiology, Development and Neuroscience, University of Cambridge, Cambridge, CB2 3EG, UK; Broad Institute of Massachusetts Institute of Technology and Harvard, Cambridge, MA 02142, USA.

**Keywords:** organoid, EGF, FGF, Integrin, KTR reporter

## Abstract

Multipotent epithelial progenitor cells can be expanded from human embryonic lungs as organoids. and maintained in a self-renewing state using a defined medium. The organoid cells are columnar, resembling the cell morphology of the developing lung tip epithelium *in vivo*. Cell shape dynamics and fate are tightly coordinated during development. We therefore used the organoid system to identify signalling pathways that maintain the columnar shape of human lung tip progenitors. We found that EGF, FGF7 and FGF10 have distinct functions in lung tip progenitors. FGF7 activates MAPK/ERK and PI3K/AKT signalling and is sufficient to promote columnar cell shape in primary tip progenitors. Inhibitor experiments show that MAPK/ERK and PI3K/AKT signalling are key downstream pathways, regulating cell proliferation, columnar cell shape and cell junctions. We identified integrin signalling as a key pathway downstream of MAPK/ERK in the tip progenitors; disrupting integrin alters polarity, cell adhesion and tight junction assembly. By contrast, stimulation with FGF10 or EGF alone is not sufficient to maintain organoid columnar cell shape. This study employs organoids to provide insight into the cellular mechanisms regulating human lung development.

**Summary statement:** RTK signalling activated MAPK/ERK and PI3K/AKT signalling regulates the shape and junctional structure of human lung epithelial progenitor cells during branching.

## INTRODUCTION

The lung undergoes branching morphogenesis to build a tree-like structure during development. In the mouse embryonic lung, a SOX9^+^ID2^+^ epithelial population located at the distal branching tips is a multipotent progenitor population, which gives rise to all lung epithelial lineages (Alanis et al., 2014; Rawlins et al., 2009). Recent work in the developing human lung has identified similar distal tip epithelial progenitors that are SOX9^+^SOX2^+^ (Danopoulos et al., 2019; Miller et al., 2018; Nikolic et al., 2017). Human epithelial progenitor cells derived from the pseudoglandular stage (∼5 to 17 post conception weeks, pcw) have been cultured as self-renewing organoids in the presence of EGF, FGF and Wnt activation and TGFB and BMP inhibition (Miller et al., 2018; Nikolic et al., 2017).

FGF10 plays a critical role in the emergence of the mouse lung during development (Arman et al., 1999; Min et al., 1998; Sekine et al., 1999). Both *in vivo* and *in vitro* studies have shown that FGF10 promotes mouse lung branching at the pseudoglandular stage (Abler et al., 2009; Agha et al., 2018; Bellusci et al., 1997; Taghizadeh et al., 2020; Volckaert et al., 2013; Weaver et al., 2000; Yin and Ornitz, 2020). Mesenchymal FGF10 acts via epithelial FGFR2, which activates MAPK/ERK signalling and promotes SOX9 expression and morphogenesis (Abler et al., 2009; Chang et al., 2013; Tang et al., 2011; Yin and Ornitz, 2020). However, recent studies on developing human lungs have not supported such a crucial role for FGF10 (Danopoulos et al., 2019; Nikolic et al., 2017). In the mouse, FGF7 deficiency does not result in severe lung defects (Guo et al., 1996) and its role in the development is thought to be secondary (Post et al., 1996; Scott et al., 1995). However, FGF7 is a contributing component to sustaining the *in vitro* culture of human tip epithelial organoids (Miller et al., 2018; Nikolic et al., 2017) and is therefore considered to contribute to tip progenitor maintenance in vivo. FGF ligands are widely expressed in the developing human lung (He et al., 2022) and neither FGF7 nor FGF10 shows regionalized distribution (Danopoulos et al., 2019).

EGFR is also implicated in lung development. *Egfr* knock-out mice show defects in lung branching and alveolar formation (Kheradmand et al., 2002; Miettinen et al., 1997). EGFR is expressed in human embryonic lungs undergoing alveolar differentiation (Klein et al., 1995). However, the function of EGF and EGFR in human lung development remains largely unknown.

We find that EGF, FGF7 and FGF10 have distinct functions in human lung tip progenitor cell shape maintenance. FGF7 promotes columnar cell shape in primary tip progenitor cells, whereas EGF and FGF10 cannot. Only FGF7 induces sustained MAPK/ERK and PI3K/AKT pathway activity when assayed using kinase translocation reporters. Moreover, inhibition experiments show that both downstream pathways are required for cell shape maintenance. The MAPK/ERK and PI3K/AKT pathways regulate proliferation, promote columnar cell shape and maintain junction organisation. We show that integrin signalling is a key pathway downstream of MAPK/ERK in the tip progenitor cells. Disrupting integrin alters cell polarity, cell adhesion and tight junction assembly. This work provides a convenient platform for dissecting RTK ligand function in epithelial cells and will facilitate future studies of cell shape and cell fate coordination in organoid-based research.

## RESULTS

### RTK signalling plays a key role in maintaining human lung tip epithelial cells as organoids

We characterised the cell shape of tip and stalk epithelial cells in the developing human lung at the early pseudoglandular stage (7 to 13 pcw) and compared *in vivo* cell shape and arrangement to the tip cells expanded *in vitro* as self-renewing (SN) organoids. Tip progenitor cells (SOX9^+^) *in vivo*, and cells in an SN organoid, were columnar and apically constricted (Fig. 1A, B; S1A). Both in *vivo* tip progenitors and SN organoids are highly proliferative and contain rounded (proliferating) cells at the basal side (Fig. 1B; S1A, arrow heads). A 3D reconstructed SN organoid is partially reminiscent of the conformation of an extending tip *in vivo* (Video S1). By contrast, *in vivo* stalk epithelial cells (SOX9^-^, adjacent to the tip), were more cuboidal and arranged into a single layer (Fig. 1A). We quantified lateral, apical and basal lengths of the tip and stalk cells and organoids (Fig. 1C; S1B-E) showing an overall conservation of tip cell shape *in vitro*, particularly of lateral cell length.

**Figure 1.**
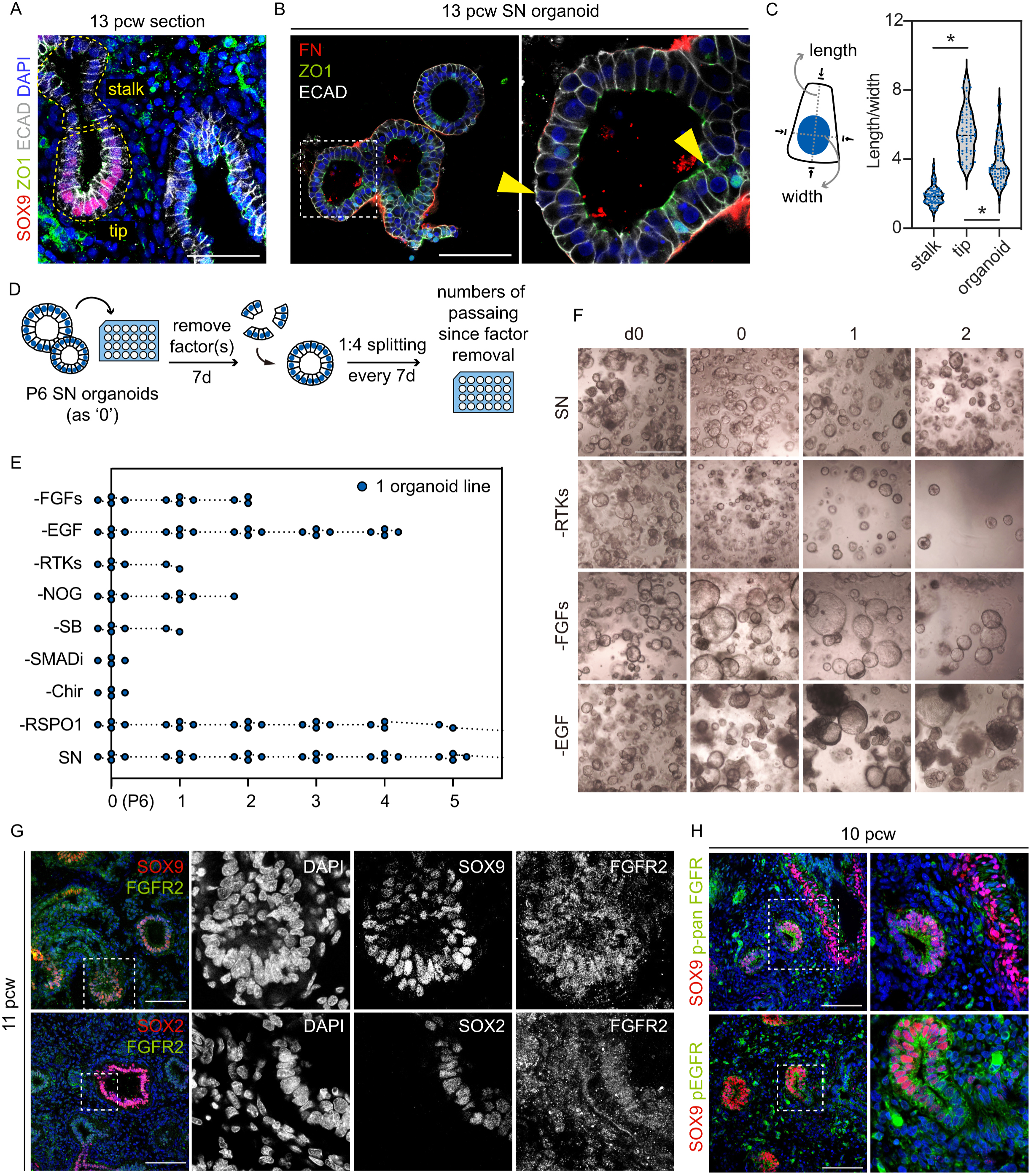
RTK signalling plays a key role in maintaining human lung tip epithelial cells. (A) Localization of ZO1 and E-cadherin (ECAD) in a 13 pcw human lung. The distal tip progenitor cells (SOX9^+^) are columnar and the stalk epithelial cells (SOX9^-^) are more cuboidal. (B) ZO1, E-cadherin and fibronectin (FN) outline the cell shape in the SN organoids. Yellow arrowheads indicate examples of cells not included in cell shape quantitation in (C). (C) Measurement of length and width of a cell through the centre of the nucleus (left) and quantitation of tip and stalk epithelial cells *in vivo*, and SN organoids. Mean±s.e.m. are shown. Blue dots show individual measurements. \**P*<0.05 (Mann-Whitney U test, N = 3 biological replicates). (D) Experimental design: P6 SN organoids were cultured in media lacking one or more components from the SN medium. Surviving organoids were passaged every 7d. (E) Survival plot of organoids cultured in each condition. N = 4 organoid lines (6 to 9 pcw) were tested (each point represents one surviving organoid line). (F) Representative images showing organoids cultured in different conditions (-FGFs: remove FGF7 and FGF10, -RTKs: remove FGF7, FGF10 and EGF, -SMADi: remove Noggin and SB431542). (G) Expression pattern of FGFR2 (detecting C-terminal cytoplasmic domain of FGFR2) in 11 pcw human lungs (2 biological replicates). (H) Expression of phospho-pan-FGFR (upper) and phospho-EGFR (lower) in a 10 pcw lung. Scale bars = 100 μm (A, B, G, H); 1mm (F).

The seven-factor organoid self-renewing (SN) medium promotes proliferation, columnar cell morphology and organoid budding (Nikolic et al., 2017). To identify the specific components responsible for some of these effects, we removed SN medium components and examined organoid survival and morphology during continued passaging (Fig. 1D). When the Wnt agonist Chir99021 (Chir), or the two SMAD inhibitors Noggin and SB431542 (SB), were removed, organoid growth terminated within seven days (Fig. 1E). By contrast, following removal of all RTK ligands (EGF, FGF7, FGF10) organoids survived for slightly longer, suggesting that RTK signalling is not required for immediate cell survival (Fig. 1E,F). Removal of EGF alone, or FGF7/10, permitted organoid culture for slightly longer than removal of all three RTK ligands, although neither media could sustain long-term culture with regular passaging (Fig. 1E,F). We noticed that organoids gradually became more spherical when FGF cytokines were removed, whereas removing EGF did not cause a striking morphology change (Fig. 1F). These data suggest a requirement for FGFR2 and EGFR signalling in the long-term maintenance of the SN organoids. We therefore explored the spatial distribution of FGFR and EGFR receptors *in vivo*.

In agreement with previous findings (Danopoulos et al., 2019), in the developing human lung at the early pseudoglandular stage (7 to 13 pcw) FGFR2 is expressed throughout the tip, stalk and airway epithelium (Fig. 1G, S1F). Likewise, EGFR localized to tip, stalk and airway epithelial cells and many mesenchymal cells (Fig. S1G). We further examined the expression of phospho-pan-FGFR and phospho-EGFR and observed subsets of cells that were responding to RTK signals throughout the epithelial compartment (Fig. 1H; S1H). These data confirm that human lung distal tip epithelium is responding to FGF and EGF signalling *in vivo* and we continued to explore the roles of these signals using the organoid model.

### FGF7 is sufficient to induce organoid budding in primary human lung tip epithelial cells

To confirm that the primary human lung epithelial tip progenitors respond to RTK signalling, we micro-dissected human embryonic tip tissues and enriched for the epithelial cells using magnetic-activated cell sorting (MACS) selection (Fig. S2A). The EpCAM^+^ single cells organized into both spherical and budding organoids when provided with SN medium (Fig. S2A,B), and these were morphologically similar to the SN organoids previously established by plating whole tips (Nikolic et al., 2017). Similarly, the single tip cells were able to form small spheres in a basal medium containing Chir, RSPO, Noggin and SB only (Fig. S2A,B). These basal spheres could not be maintained long-term in the basal medium (Fig. S2C). However, in response to the addition of RTK ligands the basal spheres began to proliferate more quickly and underwent morphological changes (Fig. S2D,E).

Based on the plasticity of the basal spheres to respond to RTK ligands, we designed an experiment to identify the effects of individual RTK ligands on tip progenitor cells (Fig. 2A). We tested EGF, FGF7 and FGF10, the three RTK ligands in the SN medium, and FGF9 which is highly enriched in the tip epithelium *in vivo* (Nikolic et al., 2017). Spheres maintained in the basal medium remained small and spherical and did not maintain SOX9 expression (Fig. 2B- G; S2F), consistent with SOX9 being downstream of FGFR signalling in the branching mouse lung (Chang et al., 2013). Organoids supplied with EGF became bigger, but remained spherical (Fig. 2B-D; S2F). Cells in EGF were SOX9-positive and more proliferative than those in the basal medium (Figure 2F-H, S2F). Both FGF7 and FGF9 were sufficient to promote organoid budding and increase proliferation (Fig. 2B-C, S2F). Organoids in these two conditions were largely SOX9 positive, although we occasionally observed FGF7-treated organoids with patchy SOX9 expression (Fig. 2F,H, S2F,G). FGF7- and FGF9-treated cells also displayed columnar morphology, in contrast to the cuboidal-like cells in the basal spheres (Fig. S2H).

**Figure 2.**
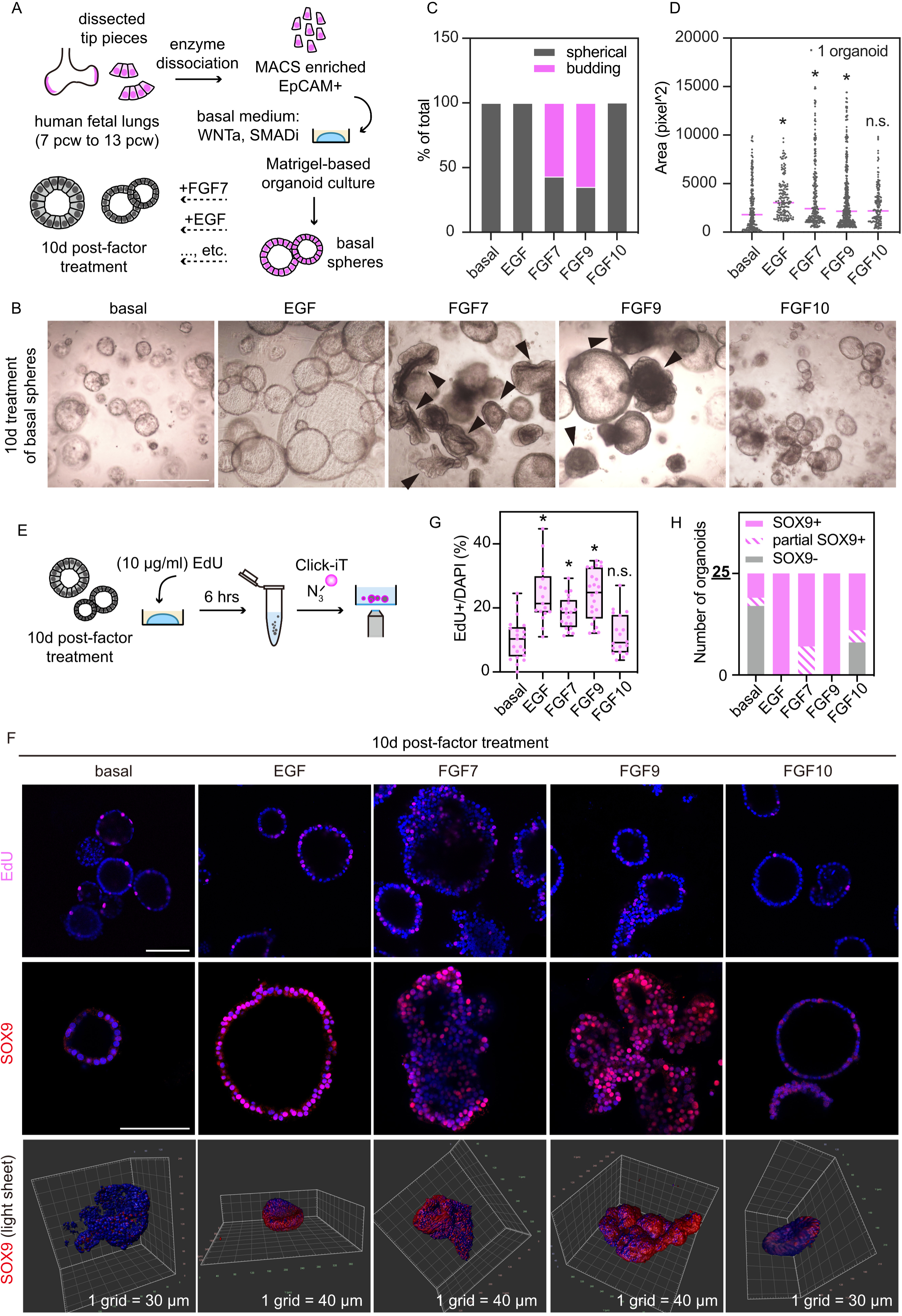
FGF7 is sufficient to induce organoid budding in primary human lung tip epithelial cells. (A) Experimental design: freshly-dissected lung tips were dissociated into single cells. Magnetic activated cell sorting (MACS)-enriched epithelial cells (5,000 per well) were seeded in Matrigel and cultured in a basal medium (RSPO1, Chir99021, Noggin, SB431542) for 10 to 14 days and grew into basal spheres. The basal spheres were distributed to different wells and supplied with specific RTK ligands for 10 days. (B) Representative images showing organoid morphologies after 10 days of RTK ligand stimulation. Black arrowheads indicate organoids scored as budding. (C) Quantitation of budding versus spherical organoids in each condition at d10, showing pooled results of 4 biological replicates. (D) Quantitation of projected area of d10 organoids by in-house ImageJ plug-in (Dr. Richard Butler), pooled results over 4 biological replicates. Means are shown. Grey dots show individual measurements. \**P*<0.05 (student’s *t* test), (E) Experimental design: after 10 days of ligand stimulation, 10 μg/ml EdU was added to each condition and incubated for 6 hours before the Click-iT assay. (F) Representative images of EdU positive cells in d10 organoids (upper). Expression pattern of SOX9 in the organoids (middle: confocal images, lower: Arivis Vision4D-reconstructed light sheet images). (G) Quantitation of EdU positive cells of d10 organoids over 4 biological replicates. ImageJ plug-in OAK was used to score the number EdU positive cells. Mean±s.e.m. are shown. Magenta dots show individual measurements. \**P*<0.05 (Mann-Whitney U test). (H) Quantitation of SOX9 expression of d10 organoids. Twenty-five organoids from 4 biological replicates were scored in each condition. Scale bars = 1 mm (B); 100 μm (F); defined on light-sheet images.

FGF10 did not have obvious effect on the basal spheres in terms of organoid morphology, proliferation, or cell shape, although it can induce some SOX9 expression (Fig. 2B-H; S2F,H). These results suggest that, unlike its crucial function in the developing mouse lung (Abler et al., 2009; Bellusci et al., 1997; el Agha et al., 2014; Li et al., 2018; Ramasamy et al., 2007; Volckaert et al., 2013), FGF10 does not work as the most critical RTK ligand in the developing human lung epithelial cells, consistent with previous publications (Danopoulos et al., 2019; Miller et al., 2018). Collectively, the results of the basal sphere stimulation experiment show that in the presence of WNT activators (Chir and RSPO), RTK signalling is an upstream regulatory pathway promoting SOX9 expression and contributing to the proliferation of the human lung tip epithelial progenitors, consistent with mouse data (Chang et al., 2013).

### FGF7 promotes columnar cell shape in primary human lung tip epithelial cells

Given the distinct organoid morphologies we observed in FGF7- and EGF-treated organoids (Fig. 2B, 2F), we focused on these two conditions to investigate the effects on tip epithelial cells in detail. Significant organoid budding in response to FGF7 treatment, or the size increase following EGF treatment occurred between days 4 and 6 (Fig. S3A). We therefore investigated organoid proliferation and cell shape at days 5 and 10. We found that proliferation of cells in FGF7 and EGF treatments differed at d5, whereas differences were no longer observable at d10 (Fig. S3B,C). Together with the observation *in vivo* that tip epithelial cells were more proliferative than stalk epithelium (Fig. S3D,E), we concluded that a high level of proliferation was required for branching *in vivo* and organoid budding *in vitro.* However, comparing basal spheres and EGF-treated organoids shows that proliferation alone is not sufficient to drive organoid budding.

To better understand the 3D volume of organoids and to inspect whether proliferating cells have a ‘preferred’ distribution or localization in the organoids, we took advantage of light sheet microscopy and carefully examined organoids of distinct architectures. Reconstructed images illustrated the complex, and mostly irregular, structures of FGF7-treated organoids and spherical, or doughnut-like, organoids in the basal medium and EGF treatment (Fig. 3A). We did not spot regionalized EdU^+^ cells in any of the d10 organoids in the three medium conditions (Fig. 3A). We saw generally consistent rates of proliferation in each condition, based on quantitation at d10 in both single z planes and 3D reconstructions (Fig. 3B).

**Figure 3.**
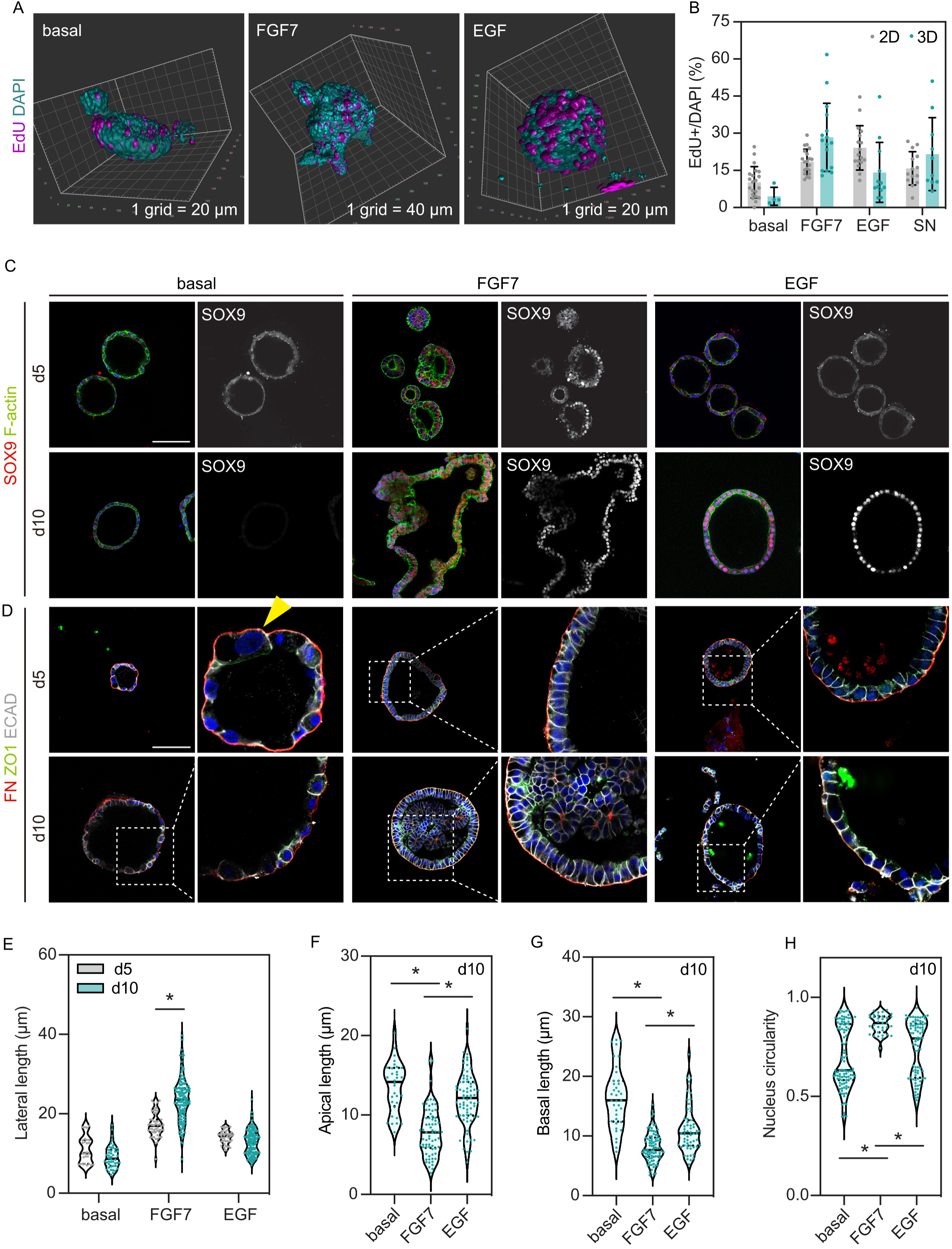
FGF7 promotes columnar cell shape in primary human lung tip epithelial cells. (A) Representative reconstructed light sheet images showing the architecture and EdU positive cells of d10 organoids. (B) Comparing quantitation of EdU positive cells of d10 organoids between confocal images (as ‘2D’) and light sheet images (as ‘3D’). Mean±s.e.m. are shown. Grey and cyan dots show individual measurements from 6 biological replicates (four replicates for 2D quantitation and three replicates for 3D quantitation with one organoid line imaged using both methods). (C) Expression pattern of SOX9 in d5 and d10 organoids. (D) Cell shape of d5 and d10 organoids. FN, fibronectin. Quantitation of cell lateral length (E), apical length (F), basal length (G) and nucleus circularity (H). Mean±s.e.m. are shown. Grey and cyan dots show individual measurements. \**P*<0.05 (Man-Whitney U test, N = 3). Scale bars = defined on light-sheet images;100 μm (C, D)

We next sought to track SOX9 expression over time. The basal spheres largely lost SOX9 expression by d5 (Fig. 3C). In contrast, FGF7-treated organoids sustained SOX9 expression over the time course (Fig. 3C). Interestingly, some organoids in the EGF treatment were not SOX9^+^ at d5, although 100% were SOX9^+^ at d10 (Fig. 3C, 2H). We conclude that both FGF7 and EGF stimulation can re-activate SOX9 expression, which is not maintained by Wnt agonists alone in the basal medium. Moreover, SOX9 re-activation likely requires a persistent high level of signalling input and therefore took several days to occur. FGF7 may have greater capacity to activate signalling pathways upstream of SOX9 than EGF.

Besides differences in promoting proliferation and sustaining SOX9 expression, FGF7- and EGF-treated organoids were distinguishable by cell shape. Cells in the basal spheres were laterally flattened with poor arrangement of ZO1 and E-cadherin, suggesting disruptions in tight and adherens junctions (Fig. 3D,E). In FGF7-treated organoids, we observed an increase in lateral cell length and an overall cuboidal-to-columnar transition over the time course (Fig. 3D,E). At d10, FGF7-treated organoids were largely reminiscent of SN organoids (Fig. 3D). By contrast, EGF treatment, although it changed cell shape, did not result in a clear cuboidal-to-columnar transition (Fig. 3D,E). Similarly, cells in the basal and EGF-treated spheres exhibited wider apical and basal surfaces than cells supplemented with FGF7 (Fig. 3C,F,G). Accordingly, we observed varied cell nucleus shape in some cells in the basal and EGF-treated spheres (Fig. 3C,H; S3F). The nuclei of these cells were wider (Fig. S3F), which might result from the lateral shortening. These data show that RTK signals tightly control the fate (SOX9 expression), cell shape and arrangement of the tip epithelial cells. These observations, that FGF7 promoted the organization of the cell junctions and cell-cell adhesion, are analogous to findings in the mouse lung where FGF10 induced genes that regulated cell adhesion (Jones et al., 2019; Lü et al., 2005).

### FGF7 efficiently activates and sustains both ERK and AKT signalling in the tip epithelial cells

The drastically different organoid morphologies and cell shapes that we observed in FGF7- and EGF-treated organoids prompted us to further investigate the downstream signalling activity. We first detailed the expression pattern of pERK and pAKT in the developing human lung (see Fig. S9D for phospho-antibody validation experiments). In the pseudoglandular mouse lungs, pERK is predominantly expressed in the distal epithelia (Hirashima and Matsuda, 2022; Jiang et al., 2018; Liu et al., 2004; Tang et al., 2011) and pAKT can be detected in the epithelial cells (Wang et al., 2005; Yin and Ornitz, 2020). However, there was sample-to-sample variation in pERK and pAKT staining in human embryonic lungs. Nonetheless, a moderately high level of pERK was observed in both the tip and the stalk epithelial cells, whereas pAKT was expressed at a reduced level in the epithelium (Fig. 4A,B, S4A,B). We did not consistently observe regionalized enrichment of pERK or pAKT between the tip and the stalk epithelia (Fig. 4A,B; S4A,B).

**Figure 4.**
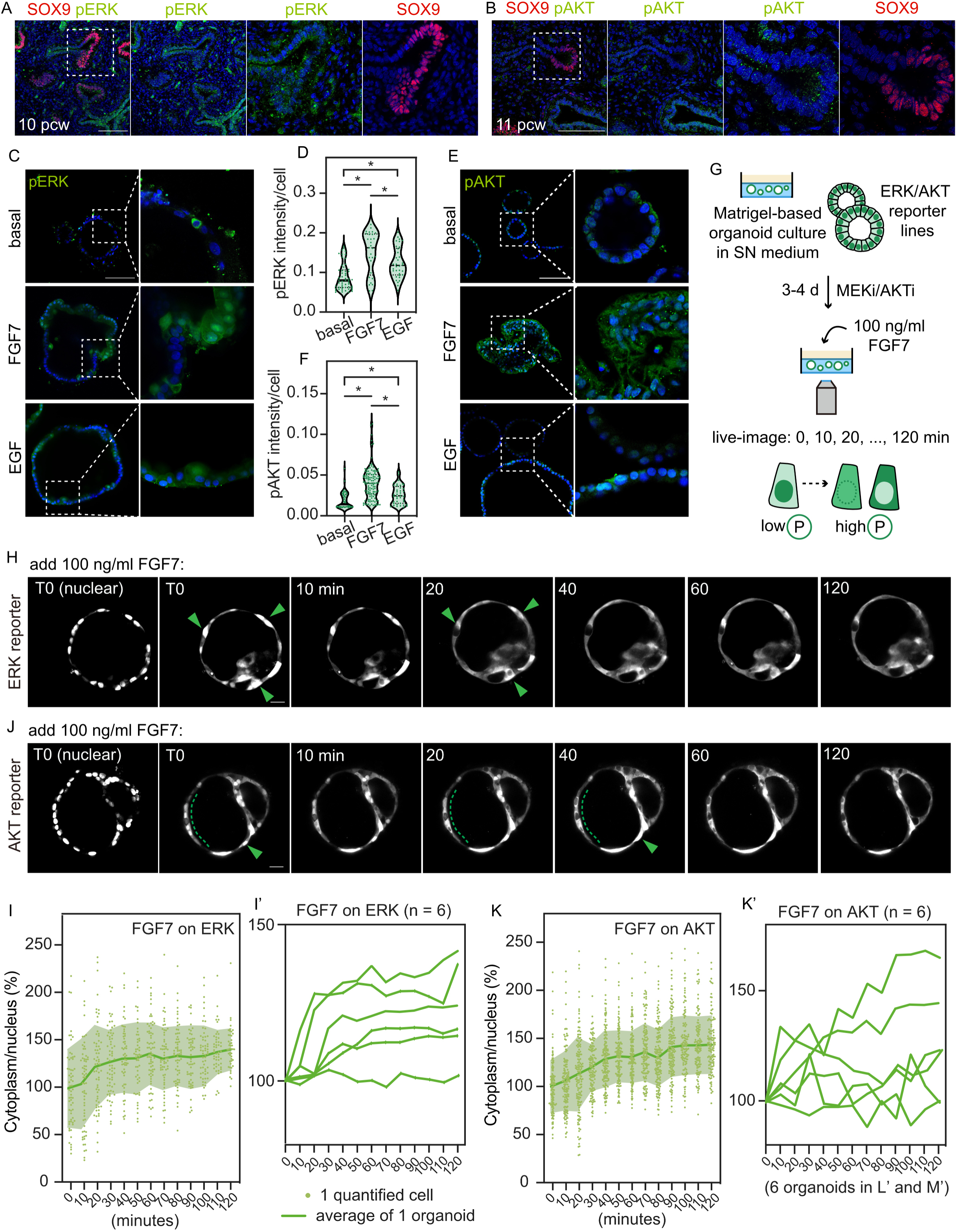
FGF7 efficiently activates and sustains ERK and AKT activation. (A,B) Expression pattern of pERK and pAKT in the early pseudoglandular lung. (C) Representative images of pERK in d10 organoids and quantitation (D), respectively. Mean ±s.e.m. are shown. Green dots show individual measurements. \**P*<0.05 (student’s *t* test, N = 3). (E) Representative images of pAKT in d10 organoids and quantitation (F), respectively. Mean ±s.e.m. are shown. Green dots show individual measurements. \**P*<0.05 (student’s *t* test, N = 3). (G) Experimental design: ERK or AKT reporter organoids were transferred into a 96-well imaging plate and cultured with MEKi (PD0325901, 200 nM) or AKTi (MK2206, 200 nM) for 3 or 4 days. 100 ng/ml FGF7 was added to the culture when live-imaging was started (T0) and the experiment lasted for 120 min. Images of multiple stacks were taken every 10 min during the live-imaging. Diagram showing the nuclear/cytoplasmic translocation of ERK or AKT reporter (right). Representative images showing the ERK reporter (H) or AKT reporter (J) cells following FGF7 stimulation. Green arrowheads and dotted lines indicate cells showing evident nucleus-to- cytoplasm fluorescence translocation. (I) Quantitation of the ERK signal obtained as the cytoplasmic/nuclear intensity for the organoid shown in H. Each point represents one scored cell, dark green line shows the average, light green line shows the 95% confidence intervals. All cytoplasmic/nuclear intensity measurements were normalized to the mean at T0 and shown as percentage. (I’) Traces of the ERK signal following FGF7 stimulation from 6 technical replicates using 3 biological replicates, shown as the average cytoplasmic/nuclear intensity per organoid per time. (K) Quantitation of the AKT signal obtained as the cytoplasmic/nuclear intensity for the organoid shown in J. Each point represents one scored cell, dark green line shows the average, light green line shows the 95% confidence intervals. All cytoplasmic/nuclear intensity were normalized to the mean at T0 and shown as percentage. (K’) Traces of the AKT signal following FGF7 stimulation from 6 technical replicates using 3 biological replicates, shown as the average cytoplasmic/nuclear intensity per organoid per time. Scale bars = 100 μm (A, B, C, E); 20 μm (H, J).

Next, we stained for the two phospho-antibodies in organoids growing in the basal medium and the FGF7, or EGF treatment and quantified the per-cell expression levels (Fig. 4C-F). The basal spheres retained ERK and AKT activation, although at much lower level than the organoids supplemented with FGF7 or EGF (Fig. 4C-F). FGF7-treated cells at d10 had pERK and pAKT staining levels greater than cells that received EGF (Fig. 4C-F). Of note, pERK and pAKT staining were heterogeneous in the FGF7- and EGF-treated organoids (Fig. 4C,E). This is expected as the staining is a snap-shot of pathway activity at 10 days after ligand treatment.

To reveal the temporal dynamics of ERK and AKT signalling in organoids subjected to different RTK ligands at a high spatiotemporal resolution, we adopted the kinase translocation reporter (KTR) system (Fig. 4G). The KTR system for ERK and AKT signalling reports on phosphorylation levels using the cytoplasmic to nuclear ratio of a fluorescent protein (Kudo et al., 2018; Maryu et al., 2016; Regot et al., 2014). We first validated the reporters, microscope set-up and image analysis by imaging the SN organoids (which have active MEK and AKT signalling) receiving a MEK or AKT inhibitor in the presence of a nuclear dye (Fig. S5A). The cytoplasm-to-nucleus fluorescence translocation indicated the reduction of ERK or AKT activity (Fig. S5B-F). Similarly, we confirmed that adding SN medium to MEK- or AKT-inhibited reporter cells could efficiently recover the cytoplasm-enriched fluorescence localization (Fig. S5G-L). These control experiments confirmed that the KTR reporters functioned as expected in our system. We next imaged dynamic changes to ERK and AKT reporters at 10 min intervals for up to 2 hours as MEK- or AKT-inhibited organoids received an FGF7 or EGF supplement from time 0 (T0, Fig. 4H,J). In these time-lapse images (Fig. 4H,J; S6, S7), we could observe that at T0 when ERK or AKT signalling was inhibited, the fluorescent reporter was located largely in the nuclei. Moreover, we were able to observe fluorescence translocation from the nuclei to the cytoplasm following ligand addition (Fig. 4H,K, arrow heads and dotted lines; S6, S7). Although there was variation between organoids, in general, FGF7 supplementation could readily initiate nucleus-to-cytoplasm translocation of both ERK and AKT reporters (Figure 4H-K, S6), but EGF could only efficiently activate ERK, but not AKT (Fig. S7). We did not notice any spatial dynamics of the ERK and AKT reporter in any of the conditions we tested. However, we could always identify more than one cell that demonstrated fluorescence translocation, which might indicate interactions between neighbouring cells. In summary, FGF7 could efficiently activate ERK and AKT signalling, whereas EGF was only effective on ERK phosphorylation.

### Activation of both ERK and AKT is required to maintain columnar cell shape and cell junctions in the tip epithelial organoids

We next asked how MAPK/ERK and PI3K/AKT signalling influenced SN organoid maintenance. We applied PD0325901 (MEK inhibitor, hereafter MEKi) or MK2206 (AKT inhibitor, hereafter AKTi) to the SN organoids to assess phenotypic changes and explore underlying molecular mechanisms (Fig. 5A; S8A). Both inhibitors displayed concentration-dependent effects on organoid budding, proliferation and cell shape (Fig. S8B-I). MEKi and AKTi application led to distinct phenotypic changes, with MEKi organoids being less proliferative and AKTi organoids having a bigger lumen (Fig. 5B, S8B,E,G). These observations suggested that the phenotypes were independently triggered by the disruption of each pathway, although the two pathways are known to interact (Rhim et al., 2016; Toulany et al., 2014). We subsequently characterized cell shape over a time course and found that the morphological differences became obvious at d4 of inhibitor treatment (Fig. 5B). Although both MEK and AKT inhibition resulted in shortened cells and disrupted tight junctions, MEKi cells were more squamous whereas AKTi cells were more cuboidal (Fig. 5B-E). At d7 the MEKi cells, which had the shortest lateral membranes, had the longest basal membranes (Fig. 5B,F). We measured nucleus circularity at d7 and found clear divergence of nucleus shape in the MEK- and AKT-inhibited organoids compared to SN cells (Fig. 5H,9A). Nuclear deformation was likely a consequence of the lateral shortening (Hatch and Hetzer, 2016; Keeling et al., 2017; Wang et al., 2022).

**Figure 5.**
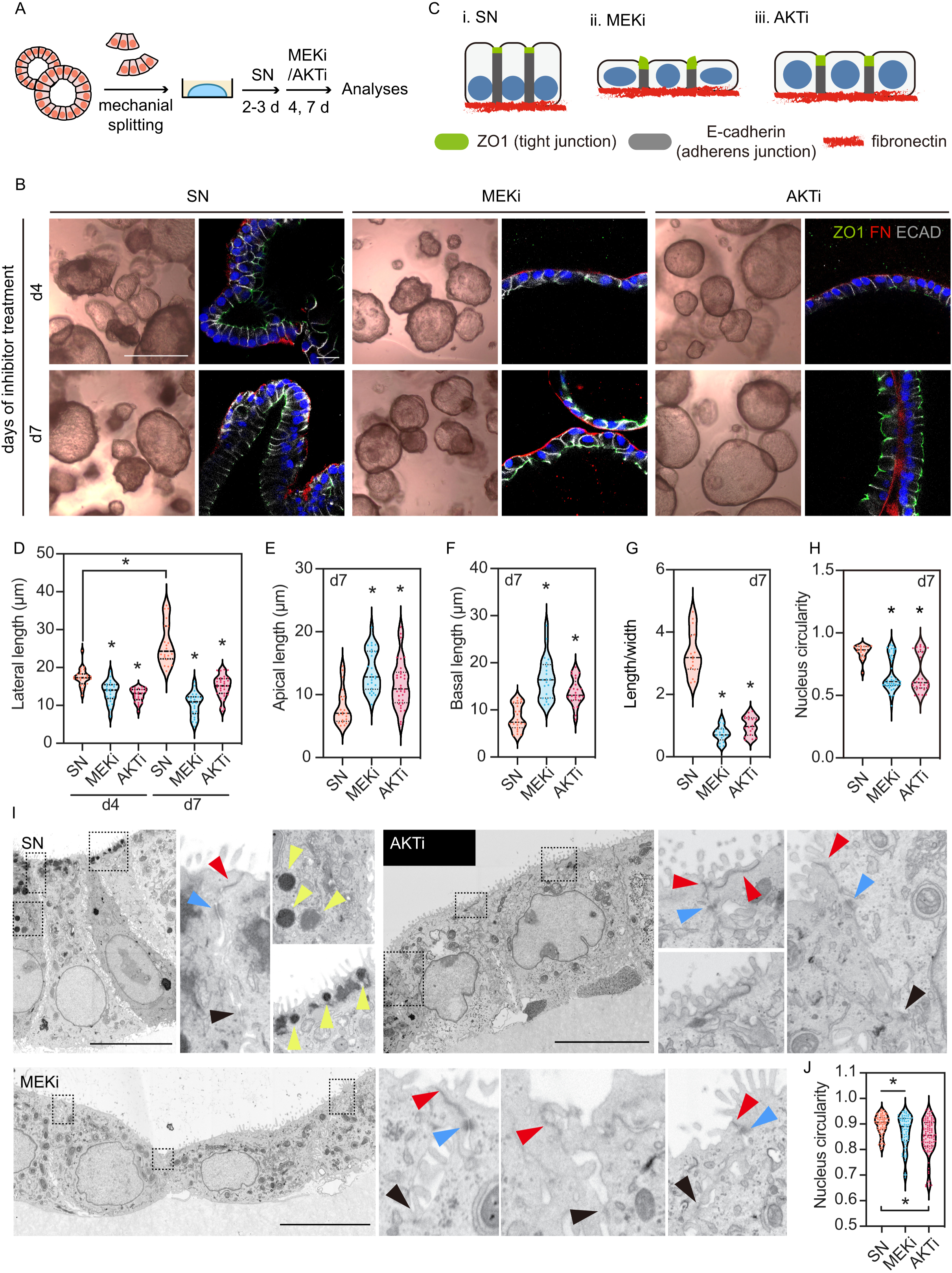
Activation of ERK and AKT is required to maintain columnar cell shape and cell junctions in the lung tip epithelial organoids. (A) Experimental design: SN organoids were treated with 200 nM MEKi (PD0325901) or 200 nM AKTi (MK2206) and analysed after 4 or 7 days. (B) Organoid morphology and cell shape of organoids at d4 and d7. Scale bar of BF image = 1 mm, scale bar of immunostaining = 20 μm. (C) Diagram showing typical cell shape of SN, MEKi and AKTi organoids. Quantitation of cell lateral length on d4 and d7 organoids (D), apical length (E), basal length (F), length/width of a cell (G) and nucleus circularity (H). Mean±s.e.m. are shown. Coloured dots show individual measurements. \**P*<0.05 (Man-Whitney U test, N = 3 biological replicates). (I) Representative SEM images showing overall cell structure, tight junction structure (zonula occluden/adheren, red arrowheads; occludin/claudin blue arrowheads), adherens junction structure (black arrowheads) and vesicles (yellow arrowheads). Scale bars = 10 μm. (J) Quantitation of nucleus circularity based on SEM images. Mean±s.d. across all scored nuclei are shown. Coloured dots show individual measurements. \**P*<0.05 (Man-Whitney U test, N = 2 biological replicates).

To investigate cell junctions at high resolution, we performed scanning electron microscopy (SEM) and observed striking differences in the SN, MEKi and AKTi cells (Fig. 5I, S9B). SN cells were overall columnar and apically constricted with basally localized nuclei that were rounded or ovoid. Microvilli and intracellular vesicles (yellow arrowheads) were clearly seen at the apical surface of the SN cells and we could identify the apical tight junctions and adherens junctions (Fig. 5I, S9B, red and blue arrowheads). In contrast, MEKi and AKTi cells were squamous or cuboidal (Fig. 5I, S9B). Although we did not notice obvious defects in microvilli in MEKi and AKTi cells, the apical surface of these cells was stretched, and the tight junction complexes were either difficult to find or mislocated (Fig. 5I, S9B). There were no apical vesicles in MEKi and AKTi cells implying perturbed protein trafficking, and nucleus shape was highly irregular (Fig. 5I,J; S9B). To summarize, our analysis of cell morphologies revealed that the two signalling pathways are required to maintain the columnar cell shape and cell junctions in the SN cells.

### Integrin genes are downstream of MAPK/ERK signalling in the human lung tip epithelial cells

We performed bulk RNA-seq to identify transcriptomic targets of PI3K/AKT and MAPK/ERK pathways in the human lung tip epithelial cells. SN organoids received 4 days of chemical inhibition prior to lysis and RNA extraction (Fig. 6A). Principal component analysis showed that inhibited cells are clearly different from the SN controls, and AKT-inhibited organoids were distinct from MEK- or ERK-inhibited (together as MAPK-inhibited, Fig. S10A). We identified > 180 differentially expressed genes (DEGs) between AKT-inhibited cells and SN cells (Fig. S10B, File. S1, log2FC > 1, *P* < 0.05). Genes that were down-regulated following AKT inhibition were highly associated with KEGG terms such as PI3K/AKT signalling, HIF- 1 signalling, and mineral absorption (Fig. S10C). By contrast, > 230 DEGs between MAPK-inhibited cells and SN control (Fig. 6B, File. S1; log2FC > 1, *P* < 0.05) were identified. We noticed the down-regulation of integrin (*ITGA2*, *ITGA6* and *ITGB4*) and cell junction-related (*ANXA10*, *JAML*, *CTNNAL1* and *GJB3)* genes (Fig. 6B). Moreover, KEGG pathway analysis revealed that genes involved in MAPK/ERK signalling, focal adhesion, actin cytoskeleton regulation and adherens junctions were down-regulated after MAPK inhibition (Fig. 6C). These data suggest that MAPK/ERK signalling regulates the cell-matrix interaction and further confirm that the signalling maintains the columnar cell shape in the human tip epithelial cells.

**Figure 6.**
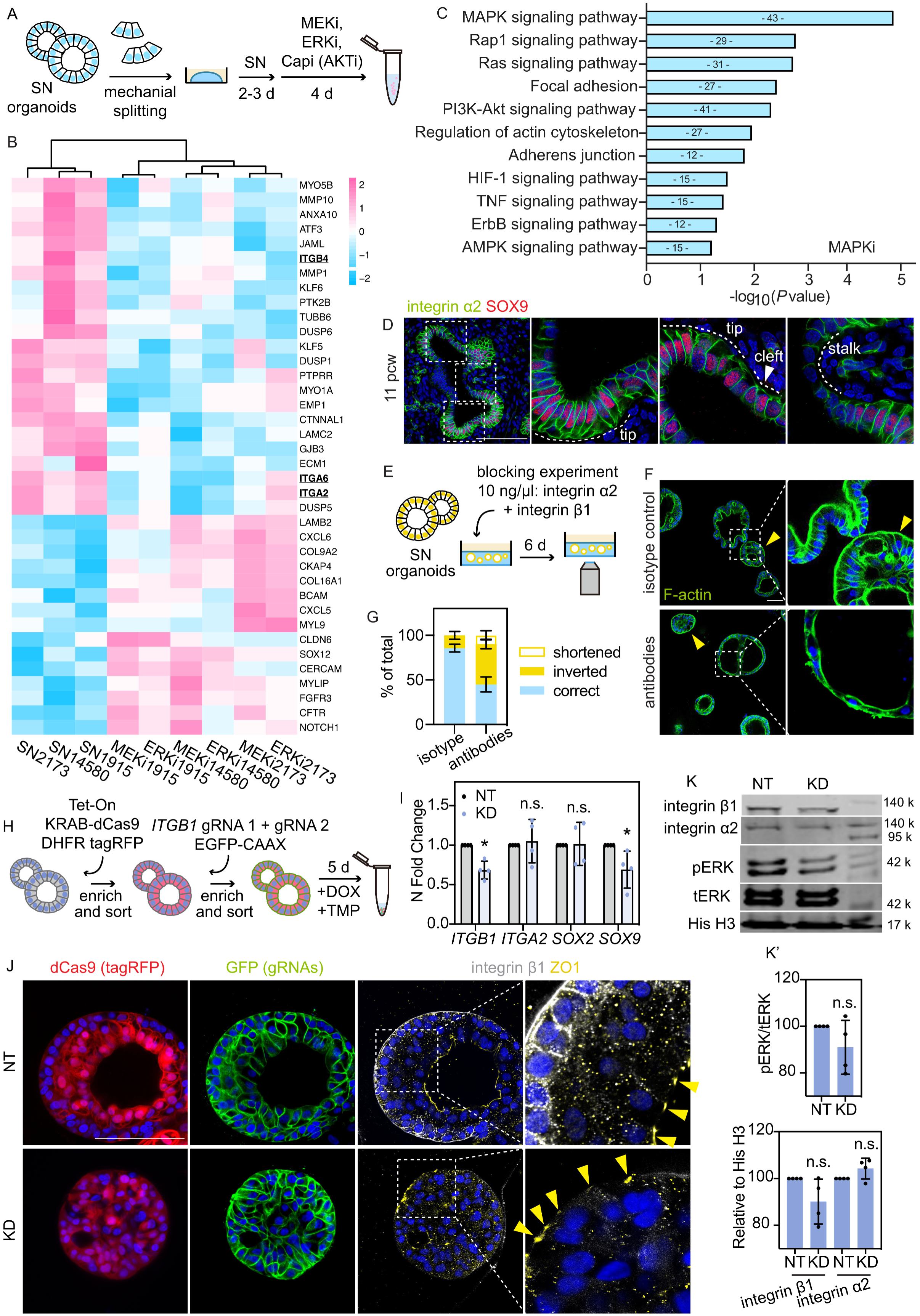
Integrin genes are downstream of MAPK/ERK signalling in the human lung tip epithelial cells. (A) Experimental design: SN organoids were treated with 200 nM MEKi (PD0325901), 200 nM ERKi (SCH772984) or 200 nM Capivasertib (‘Capi’, AKTi) and collected for RNA-seq after 4 days of treatment. (B) Heatmap showing expression level of selected genes significantly altered in MAPK- inhibited cells (N=3 biological replicates). (C) KEGG pathway analysis (selected terms) of the down-regulated genes in the MAPK- inhibited cells; log2FC > 1, adjusted p value < 0.05. (D) Localization of integrin α2 in a 11 pcw human lung. Integrin α2 was found in lateral and basal membrane of the tip and stalk epithelia with enrichment in the tips. (E) Experimental design: SN organoids were treated with 10 ng/μl integrin α2 (recombinant rabbit) and 10 ng/μl integrin β1 (mouse IgG1), or 10 ng/μl mouse IgG1 isotype control and fixed and imaged *in situ* for analysis after 6 days of treatment. (F) F-actin (ActinGreen) staining showing organoid morphology, cell shape and cell polarity of organoids treated with isotype control antibody, or integrin antibodies. Yellow arrowheads: organoids showing inverted apical-basal polarity. (G) Quantitation of organoids showing correct apical-basal polarity, inverted polarity and shortened epithelial height (with correct polarity). Mean±s.d. are shown in the bar chart, N = 3 biological replicates. (H) Experimental design: serial lentivirus transduction to generate KRAB-dCas9 expressing cells containing two gRNAs targeting *ITGB1*. Purified KRAB-dCas9^+^gRNAs^+^ cells were grown into organoids and treated with doxycycline (DOX) and trimethoprim (TMP) for 5 days. (I) qRT-PCR showing *ITGB1* downregulation by *ITGB1* gRNAs (as KD) compared to non- targeting control (as NT). Mean±s.e.m. are shown. Coloured dots show individual measurements. \**P*<0.05 (student’s *t* test, N = 4 biological replicates). (J) Representative images showing organoid morphology, cell shape and polarity of *ITGB1*- KD organoids and NT control organoids after 5 days of DOX and TMP treatment. Yellow arrowheads indicate ZO1 localization. (K) Integrin β1, integrin α2 and pERK levels of *ITGB1*-KD organoids and NT organoids after 5 days treatment of DOX and TMP. (K’) Mean and s.e.m. of integrin β1 and integrin α2 levels (normalized to histone H3) and pERK level (normalized to tERK). Black dots show individual measurements. \**P*<0.05 (student’s *t* test, N = 4 biological replicates). Scale bars = 100 μm (D, F, J).

It is well known that integrin signalling plays a key role in the interactions between epithelial cells and their ECM niches. We decided to focus especially on ITGA2, because of its *in vivo* enrichment in human lung tip epithelia, compared to the stalk, at both transcriptional and protein level, and its high abundance in organoids (Fig. 6D, File. S1) (Nikolic et al., 2017). Following MEKi treatment, immunostaining indicated that the expression pattern of integrin α2 protein was altered (Fig. S11A). Moreover, cells in basal medium-treated spheres lose lateral localization of integrin α2 (Fig. S11B), further showing that RTK inputs (FGF7, FGF10 and EGF) via MAPK/ERK activation promote integrin localisation of the human lung tip epithelial cells.

Previous studies have shown that integrins have key regulatory roles on epithelial cell shape by controlling the cytoskeleton (Domínguez-Giménez et al., 2007; Mateos et al., 2020). To test whether integrin activity is required for maintaining the columnar shape of the lung tip epithelial cells, we neutralized integrin α2 and β1 with blocking antibodies (Fig. 6E). The majority of the organoids in the isotype control condition showed correct apical-basal polarity and columnar cell shape, with some exceptions which showed inverted polarity due to organoid passaging by fragmentation (Fig. 6F, S11D, arrowhead). However, blocking antibody treatment increased the number of organoids with inverted polarity, where the cells were apically polarised towards the Matrigel and possessed more than one lumen (Fig. 6F, S11D). Meanwhile, organoids with correct polarity had laterally shortened cells, suggesting defects in adherens junctions (Fig. 6F, S11D). We therefore hypothesise that integrin signalling acts downstream of FGFR-MAPK signalling to regulate cell polarity and cell junctions in the tip epithelial cells.

To clarify the functions of integrin signalling in the tip epithelial cells, we turned to our previously published CRISPRi-based knockdown system (Sun et al., 2021) to reduce the expression of *ITGB1*, the major integrin subunit in the developing human lung (Coraux et al., 1998). SN organoids were sequentially transduced with an inducible KRAB-dCas9 vector and then a constitutive gRNA vector using lentivirus (Fig. S12), and were treated with doxycycline (DOX) and trimethoprim (TMP) to achieve gene knockdown (Fig. 6H). Experiments in four organoid lines showed a moderate, but significant and reproducible, reduction of *ITGB1* expression and a similar trend in *SOX9*, but not *ITGA2* or *SOX2* (Fig. 6I). We observed a loss of the lumen in some *ITGB1*-knockdown (KD) organoids (Fig. S12B), confirmed by immunostaining (Fig. 6J, S12C). Moreover, unlike non-targeting (NT) control organoids, the *ITGB1*-KD organoids displayed aberrant cell shape and apical-basal polarity, shown by ZO1 and integrin β1 staining (Fig. 6J, S12C). This phenotype is partially reminiscent of the integrin antibody blocked organoids (Fig. 6F, S11C). Together, these data suggest that integrin signalling indeed regulates cell polarity and cell shape of the human lung tip epithelial cells. To test whether integrin signalling is required for the tip epithelial cells to survive, we performed an organoid formation assay by seeding single control or *ITGB1*-KD cells and treating the cells with DOX and TMP for 10 days (Fig. S12D). Control cells could grow into organoids with a lumen, whereas *ITGB1*-KD cells barely re-organized into any structure (Fig. S12E).

Integrins as adhesion receptors bind to extracellular matrix proteins such as collagens and laminins to initiate ‘outside-in’ signalling and MAPK/ERK is one of the downstream pathways (Howe et al., 2002). Therefore, we examined ERK activity in d5 *ITGB1*-KD cells. Western blotting experiments demonstrated some decline in pERK level when *ITGB1* was perturbed as expected (Fig. 6K). Together, our analyses reveal that integrins are downstream of MAPK/ERK signalling in the tip epithelial cells and moreover that integrin signalling can increase ERK activity. These data implicate intricate crosstalk between integrin and MAPK/ERK signalling in the tip epithelial cells. MAPK/ERK and integrin signalling pathways both regulate the maintenance of columnar shape in the lung tip epithelial cells.

### EGF and FGF10 show combinatorial effects on the tip epithelial cells

Mesenchymal FGF10 is essential for lung morphogenesis in mice (Abler et al., 2009). However, consistent with the work of others (Danopoulos et al., 2019; Miller et al., 2018), we have shown that FGF10 addition is not sufficient to rescue columnar cell shape, proliferation, organoid budding or SOX9 expression in primary human lung tip spheres grown in basal medium (Fig. 2). Yet FGF10 is required for the establishment and maintenance of human lung bud tip organoids (Nikolic et al., 2017). We therefore sought to confirm whether FGF10 was required for the long-term maintenance of the SN organoids. We removed FGF10 from the SN medium and examined organoid survival and morphology during continued passaging (Fig. S13A-C). We noted patches of SOX2^+^SOX9^-^ cells emerging after 5 passages in <u>the</u> FGF10-removed condition (Fig. 7A, S13D,E), reminiscent of the patchy SOX9 expression in the organoids established without FGF10 (Nikolic et al., 2017). The SOX9^-^ cells in these organoids stopped proliferating (Fig. 7A), consistent with our recent findings (Sun et al., 2022) and we could not maintain these organoids for subsequent passages. Moreover, we observed altered E-cadherin patterning and F-actin in the SOX9^-^ cells (Fig. 7B, S13D), suggesting changes in cell junctions and polarity. Such observations coincided with the observations of disrupted cell junctions and laminin deposition of distal epithelial cells in *Sox* deleted embryonic mouse lungs (Rockich et al., 2013). These data confirm that FGF10 is required for long-term SN organoid maintenance where it plays a role in maintenance of progenitor cell fate, shape and proliferation.

**Figure 7.**
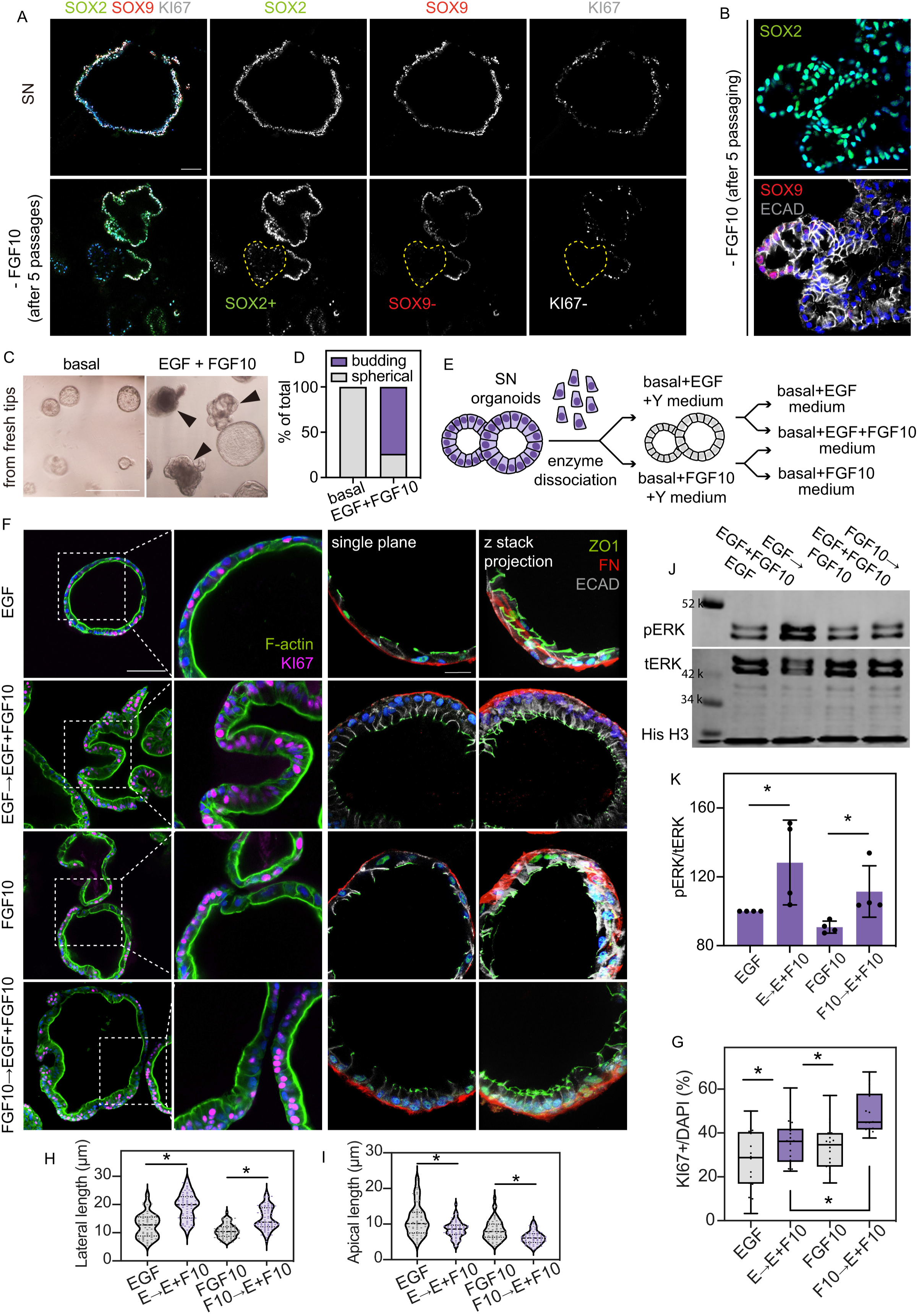
EGF and FGF10 show combinatory effects on the tip epithelial cells. (A) Representative images showing expression pattern of SOX2, SOX9 and KI67 of SN organoids and organoids in FGF10-removed medium after 5 passages. Scale bar = 100 μm. (B) Representative images showing expression pattern of E-cadherin, SOX2 and SOX9 in organoids grown in FGF10-removed medium after 5 passages. Scale bar = 50 μm. (C) Representative images showing organoid morphologies after 10 days of combined EGF and FGF10 stimulation on basal spheres cultured from freshly-dissected tip epithelial cells (compare with Fig. 2A, B). Black arrowheads show organoids scored as budding organoids. Scale bar = 1 mm. (D) Quantitation of budding versus spherical organoids in each condition at d10, pooled results of 4 biological replicates. (E) Experimental design: SN organoids were dissociated into single cells and cultured in EGF medium (basal medium plus 50 ng/ml EGF and 10 μM ROCK inhibitor, Y27632) or FGF10 medium (basal medium plus 100 ng/ml FGF10 and 10 μM ROCK inhibitor, Y27632). The EGF- and FGF10-organoids were distributed to different wells and supplied with EGF + FGF10 medium (basal medium plus 50 ng/ml EGF and 100 ng/ml FGF10) for 10 days. (F) Representative images showing organoid morphologies and proliferation (F-actin and KI67 staining) and cell shape (ZO1, FN and ECAD staining) after 10 days of combined EGF and FGF10 stimulation. Scale bar (F-actin and KI67) = 50 μm, scale bar (ZO1, FN and ECAD) = 20 μm. Quantitation of cell proliferation (G), lateral length (H) and apical length (I) on the organoids from F. Mean±s.e.m. are shown. Purple dots show individual measurements. \**P*<0.05 (Man- Whitney U test, N = 3 biological replicates). (J) pERK levels of organoids kept in the four conditions after 10 days of combined EGF + FGF10 treatment. (J’) Mean and s.e.m. of pERK level (normalized to tERK). Histone H3 was used as loading control. Purple dots show individual measurements. \**P*<0.05 (student’s *t* test, N = 4 biological replicates).

*EGF*, *FGF7* and *FGF10* are widely and ubiquitously expressed in the pseudoglandular human lungs (Danopoulos et al., 2019; He et al., 2022), meaning that the epithelial cells likely receive multiple, dynamic, RTK signalling inputs. We next interrogated the combinatorial function of FGF10 and EGF on the basal spheres (freshly dissected tip epithelial cells established in the basal medium; Fig. 2A). Interestingly, combined treatment of EGF and FGF10 robustly initiated organoid budding (Fig. 7C,D), distinct from the phenotype of single EGF or single FGF10 (Fig. 2B). To better understand such combinatorial effects, we grew single SN cells in EGF medium (basal medium plus EGF and ROCK inhibitor), or FGF10 medium (basal medium plus FGF10 and ROCK inhibitor) (Fig. 7E). In both conditions, single SN cells formed spherical organoids with cuboidal cells (Fig. 7F). Subsequently, we added EGF+FGF10 medium (basal medium plus EGF and FGF10) and observed organoid budding, cuboidal-to-columnar transition of cell shape and increased cell proliferation (Fig. 7F-I), confirming the combinatorial effect of the two ligands.

We reasoned that adding both EGF and FGF10 to the culture might result in greater downstream signalling activation than EGF or FGF10 alone, accounting for the elevated cell proliferation and columnar cell shape. Therefore, we harvested EGF or FGF10-treated organoids, and those that received both EGF and FGF10, after 10 days and examined the pERK level (Fig. 7J,K), confirming greater pERK activation following combinatorial treatment. In summary, FGF10 contributes to the maintenance of SN organoids. And we conclude that FGF7, FGF10 and EGF, the three RTK inputs in the SN medium, work together to sustain signalling activities in the tip epithelial cells to maintain progenitor identity and morphology.

## Discussion

During the early pseudoglandular stage (∼6 to 13 pcw) of human lung development, tip epithelial progenitor cells are columnar and apically constricted (Fig. 1, S1). The cells can grow into self-renewing tip epithelial organoids *in vitro* in the seven-factor SN medium (Nikolic et al., 2017). We have used the SN organoids to study the cell shape maintenance of the tip progenitor cells at the early pseudoglandular stage.

Using primary human embryonic tip epithelial cells, we showed that the RTK inputs (FGF7, FGF10 and EGF) promoted organoid budding, increased cell proliferation, and activated the ERK and AKT pathways. Among the three, FGF7 displayed the greatest potential in promoting cell proliferation and maintaining columnar cell shape. By contrast, EGF was less potent, at least partially because of its inefficiency in activating AKT, as compared to FGF7 (Fig. 4, S6, S7). Such a supportive and secondary role of EGF was previously observed in adult mouse lung epithelial organoids (Rabata et al., 2020). We initially did not observe significant effects of FGF10 on the primary human embryonic tip epithelial cells (Fig. 2), suggesting species differences between the mouse and human lung. However, we later discovered that FGF10 was required in our long-term culture, and it exerts combinatorial effects together with EGF (Fig. 7). Such observations have led us to reason that the three RTK inputs in the SN medium also show combinatorial effects on the cells. For example, our recent findings have suggested that the three RTK inputs contributed to the expression of ETV4 and ETV5, co-regulators of SOX9 in the self-renewing tip epithelial cells (Sun et al., 2022). Moreover, we speculate that similar crosstalk occurs *in vivo* where the RTK ligands cooperate and contribute to development.

Understanding the mechanisms by which cells acquire and maintain their shape is a long-standing question. Here we have shown that both ERK and AKT signalling pathways regulate the cell shape maintenance and the assembly of cell junctions in the human lung tip epithelial cells (Fig. 5). The decrease in lateral length and the increase in apical length of an SN cell is a useful proxy to describe cell shape changes and predict disruption in cell junctions. It will be interesting to broaden the cell shape examinations to later human lung developmental stages in the future and deepen the investigation of the underlying molecular processes.

We identified disruptions in the expression of integrin genes as the result of chemical inhibition of MAPK/ERK signalling (Fig. 6). In the mouse lung, several integrin genes have been identified to be implicated in the early development (Chen and Krasnow, 2012; de Arcangelis et al., 1999; Kreidberg et al., 1996; Wu and Santoro, 1996), and epithelial integrin signalling is required in the mouse lung to ensure branching morphogenesis at the pseudoglandular stage (Chen and Krasnow, 2012; Plosa et al., 2014). Our results provide experimental evidence which clarifies the roles of epithelial integrin signalling in the maintenance of cell shape and cell polarization (Fig. 6). Future investigation of the crosstalk between RTK and integrin signalling pathways would help further clarify how the niche influences the tip epithelial cells.

Taken together, our results show that RTK signalling activated MAPK/ERK and PI3K/AKT signalling regulate the shape and junctional structure of the human lung epithelial progenitor cells at the early pseudoglandular stage.

## MATERIALS AND METHODS

### Human embryonic and foetal lung tissue

Human embryonic and foetal lungs were collected from terminations of pregnancy from Cambridge University Hospitals NHS Foundation Trust under permission from NHS Research Ethical Committee (96/085) and the Joint MRC/Wellcome Trust Human Developmental Biology Resource (London and Newcastle, University College London (UCL) site REC reference: 18/LO/0822; Newcastle site REC reference: 18/NE/0290; Project 200454; www.hdbr.org). Samples used in this study had no known genetic abnormalities.

### Derivation and maintenance of human embryonic lung organoid culture

Human embryonic lung organoids were derived and maintained as previously reported (Nikolic et al., 2017). Briefly, human embryonic lung tissues were incubated in dispase (8 U/ml, ThermoFisher Scientific, 17105041) at room temperature (RT) for 2 min for dissociation. Mesenchyme was dissected away using forceps. Epithelial tips were micro-dissected and transferred into 40 μl of Matrigel (Corning, 356231) in one well of a 24-well low-attachment plate (Greiner, M9312-100EA). The plate was incubated for 15 min at 37°C to solidify the Matrigel and 600 μl self-renewal (SN) medium was added. SN medium consists of Advanced DMEM/F12 supplemented with 1x GlutaMax (ThermoFisher Scientific, 35050-061), 1 mM HEPES (ThermoFisher Scientific, 15630-060) and Penicillin/Streptomycin (as Adv+++), 1x N2 (ThermoFisher Scientific, 17502–048), 1x B27 (ThermoFisher Scientific, 12587–010), N-acetylcysteine (1.25 mM, Merck, A9165), EGF (50 ng/ml, PeproTech, AF-100-15), FGF10 (100 ng/ml, PeproTech, 100-26), FGF7 (100 ng/ml, PeproTech, 100-19), Noggin (100 ng/ml, PeproTech, 120-10C), R-spondin (5% v/v, Stem Cell Institute, University of Cambridge), CHIR99021 (3 μM, Stem Cell Institute, University of Cambridge) and SB431542 (10 μM, Bio-Techne, 1614). Once formed, SN organoids were maintained in the SN medium and passaged by mechanically breaking using P1000 pipettes every 7-10 days.

### Isolation of primary tip epithelial cells for organoid culture from single cell

The micro-dissected epithelial tips were incubated in TrypLE Express Enzyme (Thermo Fisher Scientific, 12605010) for 10 min at 37°C to dissociate into single cells. After rinsing in cold Adv+++, cells were filtered by 40 μm cell strainer and collected by centrifuge. A CD326 MicroBeads kit (Miltenyl Biotec, 130-061-101) was used to enrich EpCAM+ cells. The cell pellet was resuspended in 300 μl magnetic-activated cell sorting (MACS) buffer (0.5% BSA, 2 mM EDTA in PBS) and purified following the manufacturer’s protocol. Briefly, Fc receptors were blocked using the reagent provided to saturate non-epithelial cells, then cells were incubated with CD326 MicroBeads for 30 min at 4℃. After two MACS buffer washes, the positive cells were harvested through the LS column in the magnetic field in a 15 ml tube and transferred into 40 μl of Matrigel in 24-well low-attachment plate, 5,000 cells per well. The cells were supplied with SN medium plus Y27632 (10 μM, Merck, 688000) for the first 48 hours and SN medium onward, or basal medium plus 10 μM Y27632 for the first 48 hours and basal medium onward. Basal medium consists of Adv+++, 1x N2, 1x B27, N-acetylcysteine (1.25 mM), Noggin (100 ng/ml, PeproTech, 120-10C), R-spondin (5% v/v, Stem Cell Institute, University of Cambridge), CHIR99021 (3 μM, Stem Cell Institute, University of Cambridge) and SB431542 (10 μM, Bio-Techne, 1614).

### Immunostaining for human embryonic lung cryosections

Human embryonic lungs were fixed for 1 to 3 hours depending their size on ice in 4% (w/v) paraformaldehyde in 1x PBS, washed in 15, 20 and 30% (w/v) sucrose solutions in PBS for 1 hour each at room temperatures; before incubating in a 1:1 mix of optimal cutting temperature compound (OCT; Tissue-tek) : 30% sucrose overnight at 4℃. Lungs were embedded in 100% OCT and stored at -80℃ prior to sectioning. Human embryonic lung cryosections (12 μm) were rinsed with PBS and permeabilised with 0.2% Triton-X/PBS (washing solution). 5% normal donkey serum (Stratech, 017-000-121-JIR) in washing solution containing 0.5% (w/v) bovine serum albumin (BSA) was used for blocking at room temperature for 1 hour. Primary antibodies in blocking solution were incubated at 4℃ overnight. After three washes by the washing solution, secondary antibodies in 0.2% Triton-X/0.5 % BSA/PBS were incubated at 4℃ overnight. After three washes, DAPI (100 ng/ml, Sigma, D9542) was added for 30 min at RT. Samples were mounted in Fluoromount^TM^ Aqueous Mounting Medium (Sigma, F4680). Confocal z stacks of single planes were acquired using Leica SP8 at an optical resolution of 1024 x 1024 at 40x. Images were processed using ImageJ (version 2.1.0).

### Whole-mount immunostaining for human embryonic lung organoid culture

Organoids were recovered from the Matrigel using Corning Matrigel Cell Recovery Solution (Corning, 354253) and fixed with 4% (w/v) paraformaldehyde (PFA) for 20 min on ice. After washing in PBS for at least twice, organoids were transferred to CellCarrier-96 Ultra Microplate (PerkinElmer, 6055300) for staining. Permeabilization in 0.5% (v/v) Triton-X/PBS for 30 min was followed by washing in 0.5% (w/v) BSA, 0.2% Triton-X/PBS (washing solution). 5% normal donkey serum in washing solution was used for blocking for 1 hour at 4℃. Primary antibodies in blocking solution were incubated at 4℃ over two nights. After three washes, secondary antibodies in the washing solution were incubated at 4℃ overnight. After three washes, DAPI (100 ng/ml) was added to the washing solution for 30 min at 4℃. Organoids were mounted in a fructose-glycerol clearing buffer (Dekkers et al., 2019).

Organoids of the *ITGB1* inducible knockdown experiment (Fig. 6F,J; S12C) were immunostained *in situ* (organoids not recovered from the Matrigel). Cells were grown in the CellCarrier-96 Ultra Microplate or glass bottom microwell dishes (MatTek, P35G-1.5-20-C).

For fixation, pre-warmed 4% PFA was added to the dish after removing culture medium and incubation was 15 min at 37℃. Permeabilization was in 1% (v/v) Triton-X/PBS for 1 hour, and 1% (v/v) Triton-X was used in all solutions. Primary antibody incubation was over three nights and over two nights for secondary antibodies. The fructose-glycerol clearing buffer (Dekkers et al., 2019) was used for clearing at least overnight at 4℃.

Primary antibodies used:

|  | **Company** | **Cat. No.** | **Host species** | **Dilution** |
| --- | --- | --- | --- | --- |
| SOX9 | R&D Systems | AF3075 | Goat | 1:200 |
| SOX2 | Bio-Techne | AF2018 | Goat | 1:200 |
| ZO1 | ThermoFisher Scientific | 40-2200 | Rabbit | 1:200 |
| E-cadherin | ThermoFisher Scientific | 13-1900 | Rat | 1:1000 |
| Fibronectin | R&D Systems | AF1918 | sheep | 1:400 |
| FGFR2 (C-terminal) | Santa Cruz | SC-122 | rabbit | 1:200 |
| FGFR2 (amino terminal) | Cell Signaling Technologies | 23323S | rabbit | 1:200 |
| phospho-pan FGFR | Cell Signaling Technologies | 3476S | mouse | 1:100 |
| phospho-EGFR | Cell Signaling Technologies | 3777S | rabbit | 1:200 |
| phospho-ERK | Cell Signaling Technologies | 4370S | rabbit | 1:200 |
| phospho-AKT | Cell Signaling Technologies | 4060S | rabbit | 1:200 |
| integrin α2 | AbCam | ab181548 | rabbit | 1:400 |
| integrin β1 | AbCam | ab30394 | mouse | 1:400 |
| KI67 | BD | 550609 | mouse | 1:400 |
| CD31 | AbCam | ab9498 | mouse | 1:400 |
| GFP | AbCam | ab13970 | chicken | 1:200 |

Secondary antibodies used:

|  | Company | Cat. No. | Host species | Dilution |
| --- | --- | --- | --- | --- |
| Donkey α-mouse 488 | ThermoFisher Scientific | A21202 | mouse | 1:2000 |
| Donkey α-rabbit 488 | ThermoFisher Scientific | A21206 | rabbit | 1:2000 |
| Donkey α-goat 488 | ThermoFisher Scientific | A11055 | goat | 1:2000 |
| Donkey α-rat 488 | ThermoFisher Scientific | A21208 | rat | 1:2000 |
| Donkey α-chicken 488 | Jackson Immune | 703-545-155 | chicken | 1:2000 |
| Donkey α-mouse 594 | ThermoFisher Scientific | A21203 | mouse | 1:2000 |
| Donkey α-rabbit 594 | ThermoFisher Scientific | A21207 | rabbit | 1:2000 |
| Donkey α-goat 594 | ThermoFisher Scientific | A11058 | goat | 1:2000 |
| Donkey α-sheep 594 | Jackson Immune | 713-585-147 | sheep | 1:1000 |
| Donkey α-goat 647 | ThermoFisher Scientific | A21447 | goat | 1:2000 |
| Donkey α-rat 647 | Jackson Immune | 712-605-153 | rat | 1:2000 |

Confocal z stacks of single planes were acquired using Leica SP8 at an optical resolution of 1024 x 1024 at 40x objective. Images were processed using ImageJ (version 2.1.0).

### Western blotting

After complete removal of the Matrigel, organoids were lysed using RIPA buffer supplemented with 1x Halt^TM^ Protease and Phosphatase Inhibitor Cocktail (Thermo Fisher Scientific, 78440) on ice, with strong vortex every 5 min for 6 times. Samples were centrifuged for 15 min at 13,000 rpm, 4°C and supernatant was collected and the protein content was quantified with a BCA kit (Thermo Fisher Scientific, 23225). Equal amount of protein was denatured by mixing with 4x Laemmli buffer (Bio-Rad, 1610747) and incubated for 5 min at 95°C. Samples were separated by SDS-PAGE (Bio-Rad, 1610148) and electro-transferred onto PVDF membranes (Merck, Immobilon-P membrane). Membranes were blocked with 5% skim milk and then incubated with primary antibodies, rabbit histone H3 (1:5000, AbCam, ab1791), rabbit phospho-ERK1/2 (1:400, Cell Signaling Technologies, 4370S), rabbit total ERK1/2 (1:400, Cell Signaling Technologies, 4695), rabbit phospho-AKT (1:400, Cell Signaling Technologies, 4060S), rabbit pan AKT (1:400, Cell Signaling Technologies, 4691), rabbit integrin α2 (1:400, AbCam, ab181548) and mouse integrin β1 (1:400, AbCam, ab30394), overnight at 4°C. After extensive washes, membranes were incubated with IRDye conjugated secondary antibodies, donkey anti-mouse IRDye 800CW (1:5000, AbCam, ab216774) and donkey anti-rabbit IRDye 680CW (1:5000, AbCam, ab216779), overnight at 4°C. The protein bands were visualised using Li-Cor Odyssey system and quantified in ImageJ (version 2.1.0). Histone H3 was used as loading control. Integrin α2 and integrin β1 were normalized to histone H3. pERK and pAKT were normalized to total ERK and total AKT, respectively.

### Molecular cloning and plasmid construction

The ERK-KTR vector was generated based on Addgene #59138 with mClover swapped to mNeonGreen. The AKT KTR vector was generated based on the ERK-mNeonGreen vector with ERK-KTR sequence removed by MluI-HF and SpeI-HF cutting and swapped to AKT- FOXO3A-KTR. The AKT-FOXO3A-KTR sequence was sub-cloned from human cDNA according to Maryu et al., 2016 (Maryu et al., 2016).

For the *ITGBβ1* knock-down experiment, a doxycycline-inducible CRISPRi vector was used (Sun et al., 2021) to harvest KRAB-dCas9 cells. gRNA plasmid (Addgene #167936) was linearized with BbsI-HF restriction enzyme for 1 hour at 37°C and gel purified with QIAquick® Gel Extraction Kit (Qiagen, 28704). Two gRNAs targeting *ITGBβ1* (Horlbeck et al., 2016) were individually subcloned into gRNA vector as follows:

gRNA-1: 5’- GAGAGGCCCAGCGGGAGTCG - 3’

gRNA-2: 5’- GGGGAGACCGCAGGTGTCAG - 3’

Non-targeting control sequences: 5’- GCTGCATGGGGCGCGAATCA – 3’

### Lentiviral production and transduction of organoids

For the gRNA, dCas9 and KTR vectors, HEK293T cells at 70-80% confluency in 10 cm dishes were transfected with the insert construct plus 3rd generation packaging plasmids: pMD2.G (3 μg, Addgene plasmid #12259), psPAX2 (6 μg, Addgene plasmid # 12260) and pAdVAntage (3 μg, E1711, Promega) using Lipofectamine 2000 Transfection Reagent (11668019, Thermo Fisher Scientific) according to manufacturer’s protocol. Medium was changed after 16 hours. Lentivirus-containing supernatant was collected and pelleted (5 minutes at 1000 rpm) on the third day post-transfection. The supernatant was filtered through a 0.45 μm filter then concentrated.

To concentrate the lentiviral supernatant, 1 volume of Lenti-X Concentrator (TAKARA, 631231) was added to 3 volumes of supernatant and incubated at 4°C for 1 hour. The solution was centrifuged for 45 min at 4°C and the lentiviral pellet re-suspended in cold sterile PBS at 1/100 the original volume, distributed into 10 μl aliquots and stored at -80°C.

Lentiviral transduction of organoids was performed on a single cell solution of the organoids after TryPLE dissociation. 2.5 μl concentrated lentivirus was added to 50,000 single cells suspended in fresh SN medium supplemented with 10 μM Y27632 and incubate at 37°C overnight. Cells were collected and embedded in Matrigel at 50,000 per well the next morning and grown in SN medium supplemented with 10 μM Y27632 for 24 hours. Virally transduced cells were cultured for 7-10 days to grow into organoids. To isolate a pure population of transduced cells, single cells were collected by dissociating organoids with TryPLE and sorted based on fluorescence colours using Sony SH800S cell sorter.

### RNA extraction, cDNA synthesis, qRT-PCR and bulk RNA-sequencing

Organoids were collected from the Matrigel and lysed. RNA extraction was performed according to the manufacturer’s protocol (Qiagen, 74004). RNA concentrations were measured by NanoDrop. cDNA synthesis was performed using MultiScribe Reverse Transcriptase (Applied Biosystem, 4308228). The mix was left at 25 °C for 5 minutes, 50°C for 50 minutes then 15 minutes at 70 °C. cDNA was diluted 1:10 and 1 μl was used for each qPCR reaction with SYBR Green assays (PowerUp SYBR Green Master Mix, Applied Biosystem, 100029284). Relative Cp values for target genes were standardized against *GAPDH* expression. Relative gene expression was calculated using ΔΔCp method. *P*-values were obtained using an unpaired two-tailed student’s t-test with unequal variance.

Primers:

*GAPDH*-F: 5’-TGCCCTCAACGACCACTTTG-3’

*GAPDH*-R: 5’-GGGTCTCTCTCTTCCTCTTGTGCT-3’

*ITGB1*-F: 5’-TTCAAGGGCAAACGTGTGAG-3’

*ITGB1*-R: 5’-GGACACAGGATCAGGTTGGA-3’

*ITGA2*-F: 5’-AGAAAGCCGAAGTACCAACAGGAGT-3’

*ITGA2*-R: 5’-TGCAGGTAGGTCTGCTGGTTCA-3’

*SOX2*-F: 5’-TACAGCATGTCCTACTCGCAG-3’

*SOX2*-R: 5’-GAGGAAGAGGTAACCACAGGG-3’

*SOX9*-F: 5’-AGCACTGGGAACAACCCGTCT-3’

*SOX9*-R: 5’-TAGGATCATCTCGGCCATCTTCGC-3’

For bulk RNA-seq, RNA quality was validated using an Agilent 2200 Tapestation with High Sensitivity RNA Screen Tape (Agilent, 5067-5579). RNA samples were sent for Eukaryotic RNA-seq library preparation and sequencing (PE150) at Novogene (Cambridge, UK). mRNA was purified from total RNA using poly-T oligo-attached magnetic beads. After fragmentation, the first strand cDNA was synthesized using random hexamer primers, followed by the second strand cDNA synthesis. The library was checked with Qubit and real-time PCR for quantification and bioanalyzer for size distribution. Quantified libraries were pooled and sequenced on Illumina platforms and paired-end reads were generated. Raw sequencing data were pre-analysed by Novogene. Raw data (raw reads) of fastq format were firstly processed through Novogene in-house perl scripts. In this step, clean data (clean reads) were obtained by removing reads containing adapter, reads containing ploy-N and low-quality reads from raw data. Index of the reference genome was built and paired-end clean reads were aligned to the reference genome using Hisat2 (2.0.5). To count the reads numbers mapped to each gene, featureCounts (1.5.0-p3) was used and fragments per kilobase of transcript sequence per millions base pairs (FPKM) of gene was calculated based on the length of the gene and reads count mapped to this gene. Differential expression analysis of two conditions was performed using the DESeq2 R package (1.20.0). The resulting P-values were adjusted using the Benjamini and Hochberg’s approach for controlling the false discovery rate. KEGG pathway analysis were performed using DAVID (Huang et al., 2009). RNA-seq data have been deposited to GEO (GSE211308) and are publicly available as of the date of publication.

### KTR reporter experiments and image analysis

ERK-KTR and AKT-FOXO3A-KTR expressing organoids were cultured in CellCarrier-96 Ultra Microplate and imaged on a Zeiss 880 Airyscan inverted confocal microscope equipped with an incubation chamber and CO2 supply to maintain 37°C and 5% CO2. NucRed Live 647 (ThermoFisher Scientific, 2146834) was added 1 hour before live-imaging experiments to allow cell nucleus imaging. Multiple z stacks (5 μm step) at each time point of the KTR-mNeonGreen and the nuclear marker were simultaneously captured through a 25x0.8 N.A. water objective.

ImageJ and CellProfiler (Stirling et al., 2021) were used to process the images. To determine the nuclear and cytoplasmic fluorescence intensities shown in Fig. 4 and S5-S7, we referred to previous reports with custom changes (Kudo et al., 2018; Regot et al., 2014). We manually selected at least three non-consecutive z planes at each time point (same relative z position) in ImageJ. In CellProfiler, the nuclear region of each cell was segmented based on the NucRed 647 channel. The nuclear segmentation was used as a mask and a ring of 5 pixels width around the nucleus from the nuclear segmentation was used to define the cytoplasmic region. Fluorescence intensity for mNeonGreen of the nuclear region and cytoplasmic region of each cell was measured. The cytoplasmic to nuclear ratio (cytoplasm/nucleus ratio) of the KTR-mNeonGreen of each measured cell was calculated for each time point by dividing the mean cytoplasmic intensity by the mean nuclear intensity of a cell. This ratio is normalized to time 0 (as 100%) and shown as % in Fig. 4 and S5-S7.

### Electron microscopy imaging

The organoid samples were fixed in 2 % formaldehyde/2 % glutaraldehyde in 0.05 M sodium cacodylate buffer (NaCAC), pH 7.4, containing 2 mM calcium chloride (Merck, C27902) overnight at 4°C. After washing in 0.05 M NaCAC at pH 7.4, the samples were osmicated for 3 days at 4°C. After washing in deionised water (DIW), the samples were treated twice with 0.1 % (w/v) thiocarbohydrazide (Merck, 223220) in DIW for each 20 min and 1 hour at room temperature in the dark, followed by block-staining with uranyl acetate (2 % uranyl acetate in 0.05 M maleate buffer pH 5.5) for 3 days at 4°C. Then, the samples were dehydrated in a graded series of ethanol (50%/70%/95%/100%/100% dry) 100% dry acetone and 100% dry acetonitrile, three times in each for at least 5 min. Next, the samples were infiltrated with a 50:50 mixture of 100% dry acetonitrile/Quetol resin (TAAB, Q005) without BDMA (TAAB, B008) overnight, followed by 3 days in 100% Quetol without BDMA. The sample was infiltrated for 5 days in 100% Quetol resin with BDMA, exchanging the resin each day. The Quetol resin mixture is: 12 g Quetol 651, 15.7 g NSA (TAAB, N020), 5.7 g MNA (TAAB, M012) and 0.5 g BDMA. Samples were placed in embedding moulds and cured at 60°C for 3 days. Thin sections were cut using an Ultracut E ultramicrotome (Leica) and mounted on melinex plastic coverslips. The coverslips were mounted on aluminium SEM stubs using conductive carbon tabs and the edges of the slides were painted with conductive silver paint. Then, the samples were sputter coated with 30 nm carbon using a Quorum Q150 T E carbon coater and imaged in a Verios 460 scanning electron microscope (FEI, Thermo Fisher Scientific) at 4 keV accelerating voltage and 0.2 nA probe current in backscatter mode using the concentric backscatter detector in immersion mode at a working distance of 3.5-4 mm; 1,536 x 1,024 pixel resolution, 3 μs dwell time, 4 line integrations. Stitched maps were acquired using FEI MAPS software using the default stitching profile and 10% image overlap.

### Light sheet imaging

Whole-mount immunostained organoids were embedded in an embedding solution as previously described (Dekkers et al., 2019) with light-sheet glass capillaries. Briefly, 0.4 g of low-melting point agarose (Bio-Rad, 1613111) was fully dissolved in 10 ml of water and 10 ml of fructose-glycerol clearing solution added and mixed well to obtain a clear solution. A Zeiss Z1 light sheet microscope was used for imaging. Samples were imaged by placing the sample-containing capillary in the light sheet chamber filled with fructose-glycerol clearing solution and pushing down the embedded sample from the capillary to be exposed for imaging. A 20x detection objective (clearing immersion N.A. = 1.0) was used to acquire whole Z stack images with 1,024 x 1,024 frame size. Arivis Vision4D was used to process the Z planes and generate 3D-rendering images that exported as .tiff files.

### Antibody Blocking Experiment

Integrin α2 and β1 blocking experiment (Fig. 6E-G) was performed by culturing SN organoids in CellCarrier-96 Ultra Microplate. 50 μl of Matrigel containing organoids was seeded in each well. Organoids recovered from cell seeding were treated with 10 ng/ml recombinant rabbit integrin α2 (AbCam, ab181548) and 10 ng/ml mouse IgG1 integrin β1 (AbCam, ab30394), or 10 ng/ml mouse monoclonal 2C11 IgG1 isotype control antibody (AbCam, ab1927) for 6 days with culture medium and antibodies being replenished every 2 days. Organoids were fixed *in situ* and stained for F-actin (488 ReadyProbes Reagent, ThermoFisher Scientific, R37110) according to manufacturer’s instructions.

### EdU assay

Click-iT EdU Imaging Kit (ThermoFisher Scientific, C10338) was used to assay organoids shown in Fig. 2F according to manufacturer’s instructions. Briefly, 10 μg/ml EdU was added to organoids in different conditions for 6 hours. Afterwards, EdU-containing media were washed off and organoids were recovered from Matrigel and fixed with 4% PFA for Click-iT assay.

### Inducible knockdown of *ITGB1* in the organoids

SN organoids were sequentially transfected with inducible KRAB-dCas9 vector and then gRNA vector or non-targeting control vector (Fig. S12). Purified cells were cultured into organoids and treated with 2 μg/ml doxycycline (DOX, Merck, D9891) and 10 μM trimethoprim (TMP, Merck, 92131) for 5 days. Samples were then collected for qRT-PCR and western blotting, or fixed using pre-warmed 4% PFA for 15 min at 37℃ *in situ* for immunostaining.

### Image quantitation

Cell shape quantitation was performed in ImageJ (version 2.1.0). Images of cryosections and organoids were manually scored by drawing lines and measuring the lengths, based on ZO1, fibronectin and E-cadherin staining or F-actin staining. Cells not integrated into the epithelial sheet were not included, because cells round up during division.

Nucleus circularity (DAPI staining based) was measured and calculated in ImageJ (version 2.1.0) by manually tracing the outline of individual cell nuclei following their DAPI signal, or circling the nucleus of SEM images.

Quantitation of EdU assay was performed in ImageJ (version 2.1.0) with 2D images acquired by Leica SP8 microscope. The Fiji plug-in OAK (Organoid Analysis Kit available at https://github.com/gurdon-institute/OAK/releases/tag/1.7.1) was used to score the EdU positive cell numbers and total cell numbers. Similarly, KI67 quantitation was achieved with the same procedure. Arivis Vision4D was used to quantify EdU positive cells with images acquired by Zeiss Z1 light sheet microscope by scoring cell numbers from the EdU channel and DAPI channel.

Quantitation of spherical or budding organoids (Fig. 2C) was performed by manually counting organoid numbers of each phenotype.

Quantitation of the area of the organoids (Fig.2D) was performed by using a custom script for Fiji (File S1) to segment and measure organoids in images taken on Zeiss Axiophot compound microscope.

Quantitation of SOX9 expression of organoids (Fig. 2H) was performed by sampling 25 SOX9- stained organoids from three biological replicates and manually scoring the numbers of SOX9+, partially SOX9+ and SOX9- organoids.

Quantitation of pERK/pAKT intensity per cell (Fig. 3A) was performed in CellProfiler. Nuclear region of each cell was segmented based on DAPI staining. The nuclear segmentation was used as a mask and a ring of 5 pixels width around the nucleus from the nuclear segmentation was used to define the cytoplasmic region. Fluorescence intensity for pERK or pAKT of the nuclear region and cytoplasmic region of each cell was measured and added up as per cell intensity.

Quantitation for the integrin blocking experiments was performed by sampling F-actin stained images of isotype control and integrin blocking condition and count the numbers of organoids showing correct apical-basal polarity, inverted polarity or shortened cell height (but with correct polarity).

### Quantification and statistical analysis

Data are illustrated as mean ± standard deviation (SD) or mean ± standard error of the mean (s.e.m.) as stated in the figure legends. Statistical significance was evaluated by Mann-Whitney U test or unpaired Student’s *t* test; n.s.: not significant, \**P* < 0.05.

## ACKNOWLEDGEMENTS

We acknowledge the Gurdon Institute Imaging Facility and Dr Karin Mueller of Cambridge Advanced Imaging Centre for microscopy support.

For the purpose of open access, the author has applied a Creative Commons Attribution (CC-BY) licence to any Author Accepted Manuscript version arising from this submission.

## COMPETING INTERESTS

The authors declare no competing or financial interests.

## AUTHOR CONTRIBUTIONS

Conceptualization: S.L., E.L.R.; Methodology: S.L., D.S., E.L.R.; Image analysis: S.L., R.B.; Software: R.B.; Investigation: S.L.; Writing – original draft: S.L., E.L.R.; Writing – reviewing & editing: S.L., E.L.R.; Supervision: E.L.R.; Project administration: E.L.R.; Funding acquisition: E.L.R.

## FUNDING

D.S. is supported by a Wellcome Trust PhD studentship (109146/Z/15/Z) and the Department of Pathology, University of Cambridge. E.L.R is supported by the MRC (MR/P009581/1). And core funding to the Gurdon Institute from the Wellcome Trust (203144/Z/16/Z) and CRUK (C6946/A24843).

---

**Figure S1.**
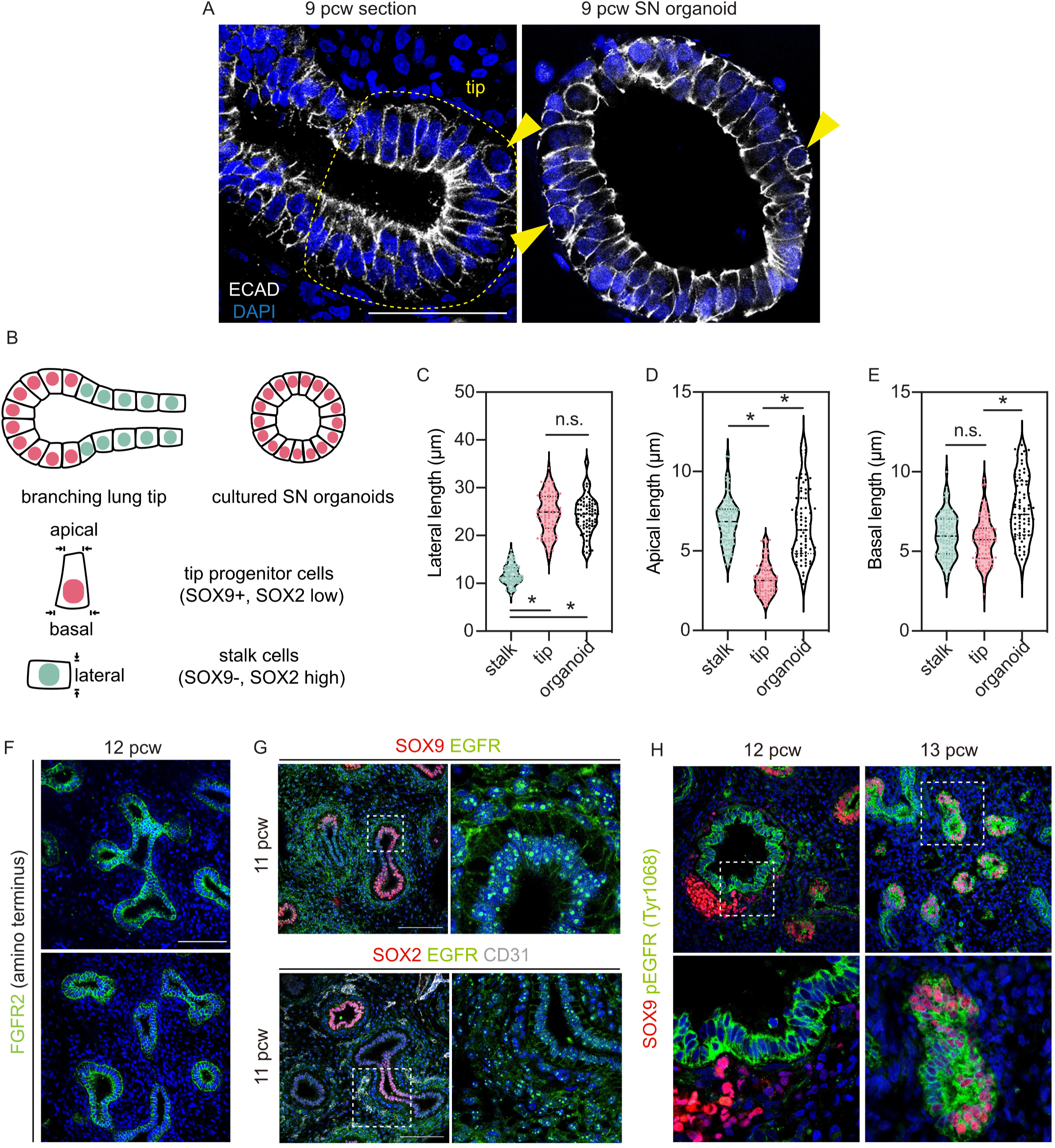
Characterization of the epithelial cell shape and receptor expression in the pseudoglandular stage human lungs. (Related to Figure 1) (A) Representative images showing the comparison of cell shape and arrangement of the tip epithelial cells *in vivo* and SN organoid cells *in vitro*. Yellow arrowheads show proliferating cells. (B) Diagram showing the quantitation strategy of the epithelial cells. Apical, basal and lateral length of a fully integrated cell were manually measured in ImageJ based on E-cadherin and ZO1 staining, or F-actin staining. Measurement of the lateral (C), apical (D) and basal (E) lengths of tip progenitor cells and stalk epithelial cells *in vivo*, and SN cells of organoids. Mean±s.e.m. are shown. Blue dots show individual measurements. \**P* < 0.05 (Man-Whitney U test, N = 3 biological replicates). (F) Expression pattern of FGFR2 (detecting amino terminus) in the early pseudoglandular lung. (G) Expression pattern of EGFR in the early pseudoglandular lung. (H) Expression pattern of phospho-EGFR in the early pseudoglandular lung. (Note that the mesenchymal SOX9 seen in the left panel is developing cartilage surrounding the more proximal airways). Scale bars = 50 μm (A); 100 μm (F, G, H).

**Figure S2.**
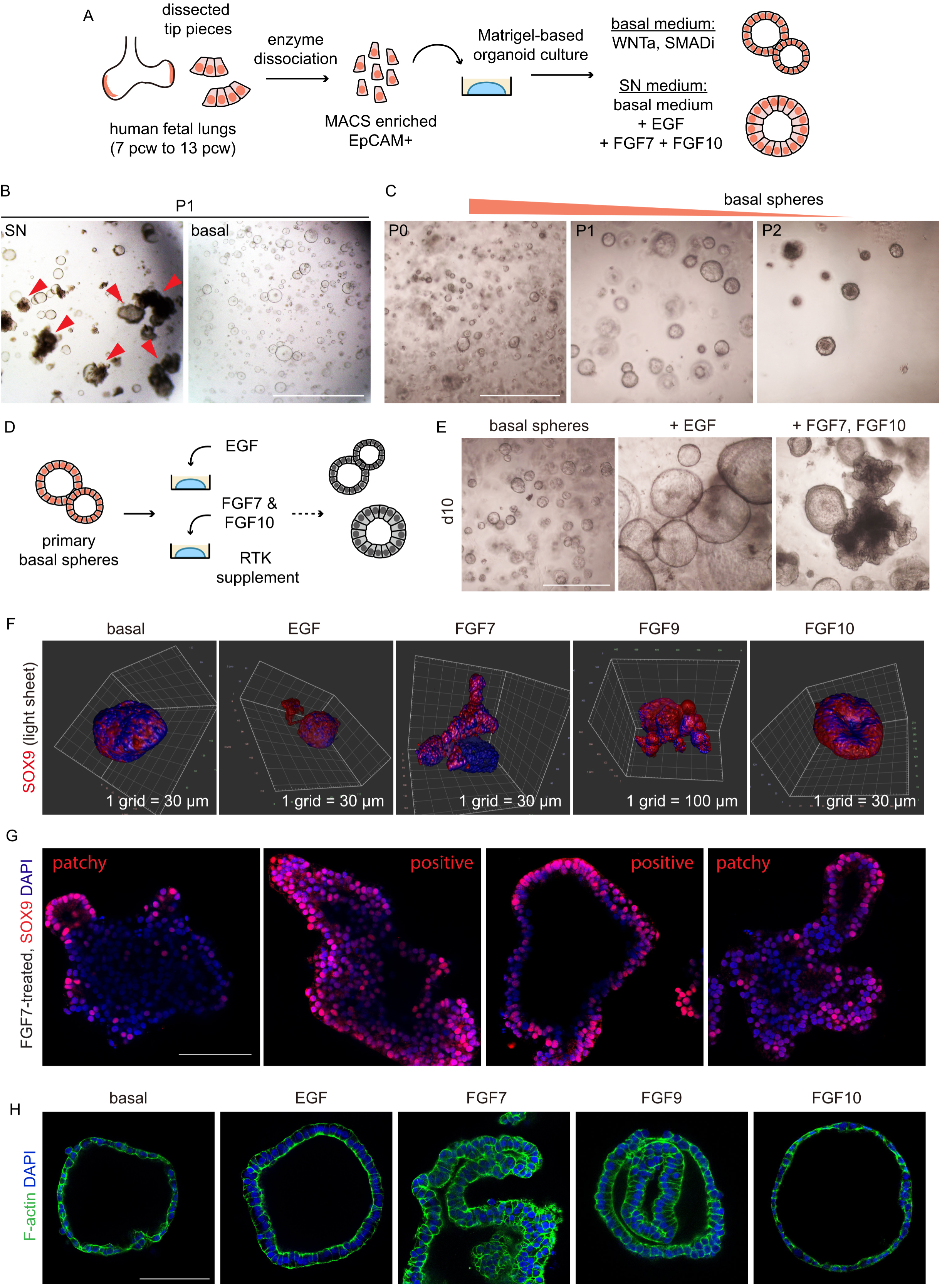
Establishment of the basal sphere culture that enabled ligand stimulation assays. (Related to Figure 2) (A) Experimental design: freshly-dissected lung tips were dissociated into single cells. MACS-enriched epithelial cells (5,000 per well) were seeded in Matrigel and cultured in a basal medium (RSPO1, Chir99021, Noggin, SB431542) or SN medium (basal medium plus EGF, FGF7 and FGF10) and grown into 3D structures. (B) Representative images showing organoid morphologies in the basal medium or SN medium. Red arrowheads showing budding organoids. (C) Representative images showing basal spheres at different passages. (D) Experimental design: primary basal spheres were distributed to different wells and received different RTK (EGF, FGF7 or FGF10) supplements for 10 to 14 days. (E) Representative images showing organoid morphologies in different conditions. (F) Light sheet images (reconstructed in Arivis Vision4D) showing expression pattern of SOX9 in the d10 organoids. Related to Figure 2F. (G) Expression pattern of SOX9 in the FGF7-treated organoids. Related to Figure 2F. (H) F-actin (ActinGreen 488) staining showing organoid morphology and cell shape of the d10 organoids. Scale bars = 1 mm (B, C, E); 100 μm (G, H).

**Figure S3.**
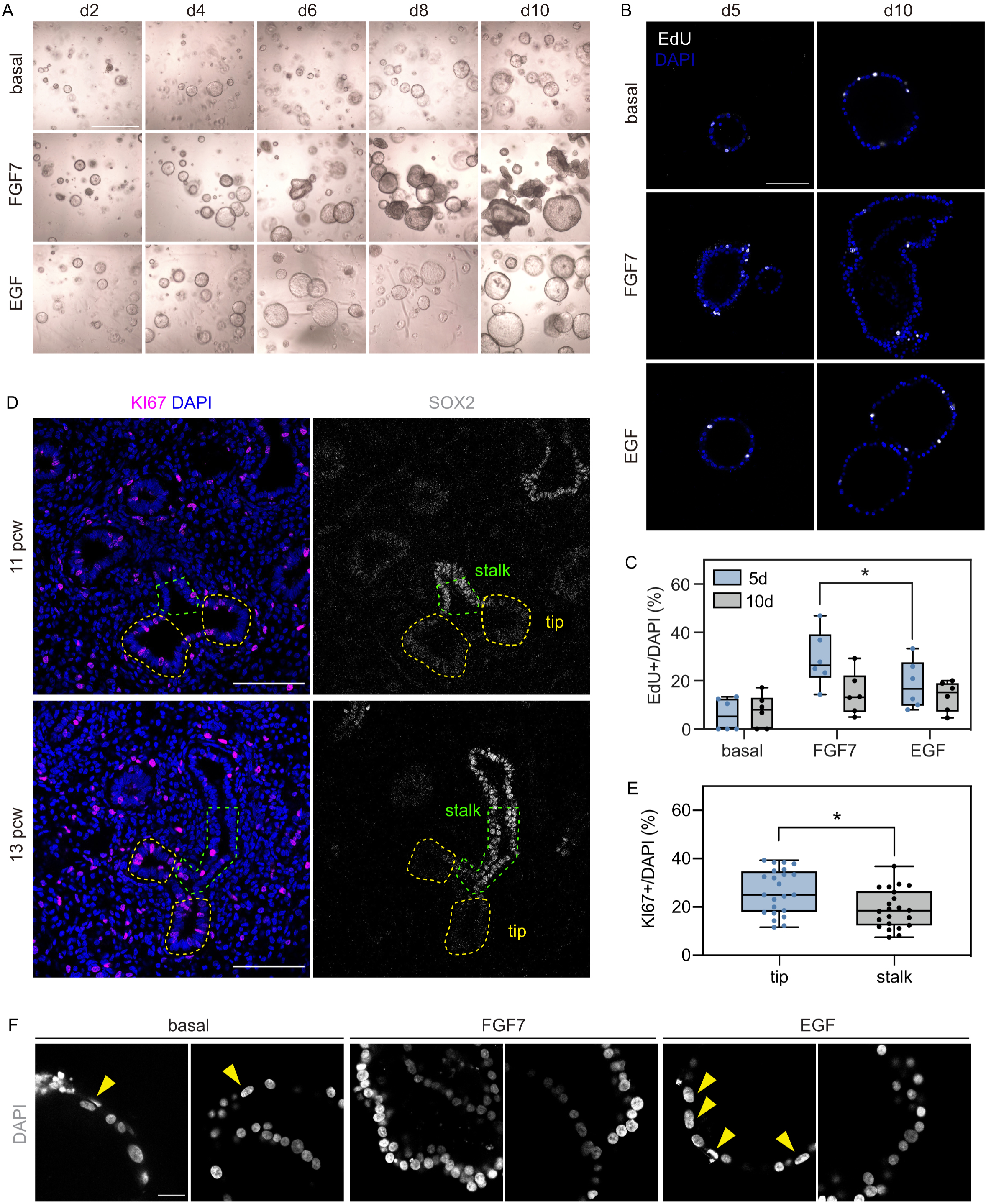
Time-course characterization of FGF7- and EGF-treated organoids. (Related to Figure 3) (A) Representative images showing the time-course organoid morphologies following FGF7 or EGF stimulation. (B) EdU assay showing proliferating cells at d5 and d10 (protocol shown in Fig. 2E). (C) Quantitation of EdU positive cells of d5 and d10 organoids. In house ImageJ plug-in OAK (Dr. Richard Butler, available at https://github.com/gurdon-institute/OAK/tree/master/OAK) was used to score the number EdU positive cells. Mean±s.e.m. are shown. Blue and black dots show individual measurements. \**P* < 0.05 (Man-Whitney U test, N = 3 biological replicates). (D) Representative images showing cell proliferation in the early pseudoglandular human lungs. (E) Quantitation of KI67+ cells in the tip and stalk epithelia. Mean±s.e.m. are shown. Blue and black dots show individual measurements. \**P* < 0.05 (Man-Whitney U test, N = 4 biological replicates). (F) Representative images showing nuclear shape of d10 organoids. Yellow arrowheads showing flat nuclei. Related to Figure 3D and 3H. Scale bars = 1 mm (A); 100 μm (B, D); 20 μm (F).

**Figure S4.**
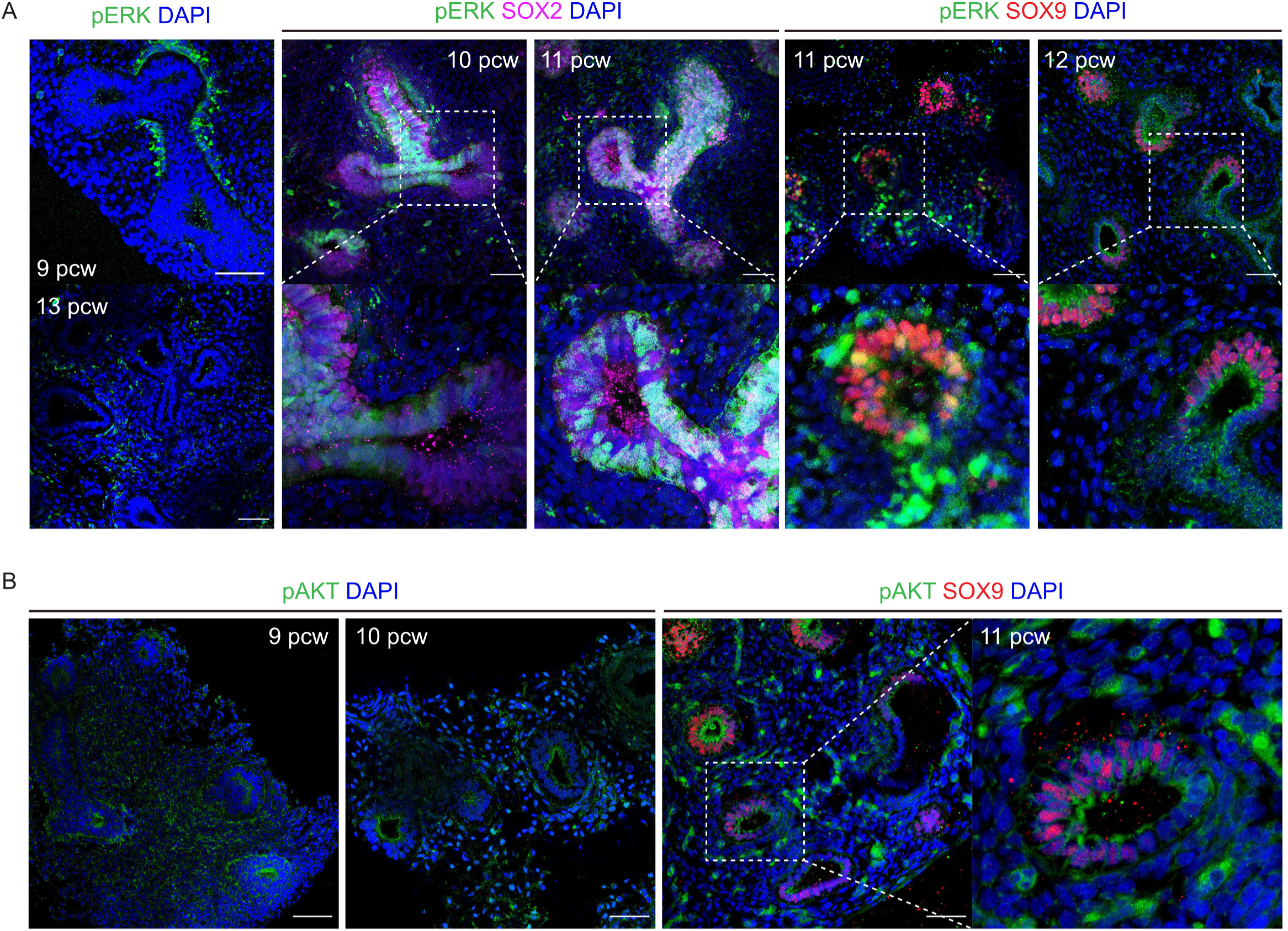
Characterization of pERK and pAKT in the pseudoglandular stage human lungs. Related to Figure 4. Expression pattern of pERK (A) and pAKT (B) in the early pseudoglandular lung. (N=6 biological replicates shown in A and 3 biological replicates in B). Note that the staining level and pattern varies extensively from sample to sample. This is probably because these phospho-proteins are turned over very rapidly and there are differences in time between isolation and fixation of each sample which is out of the control of the laboratory. Scale bars = 100 μm.

**Figure S5.**
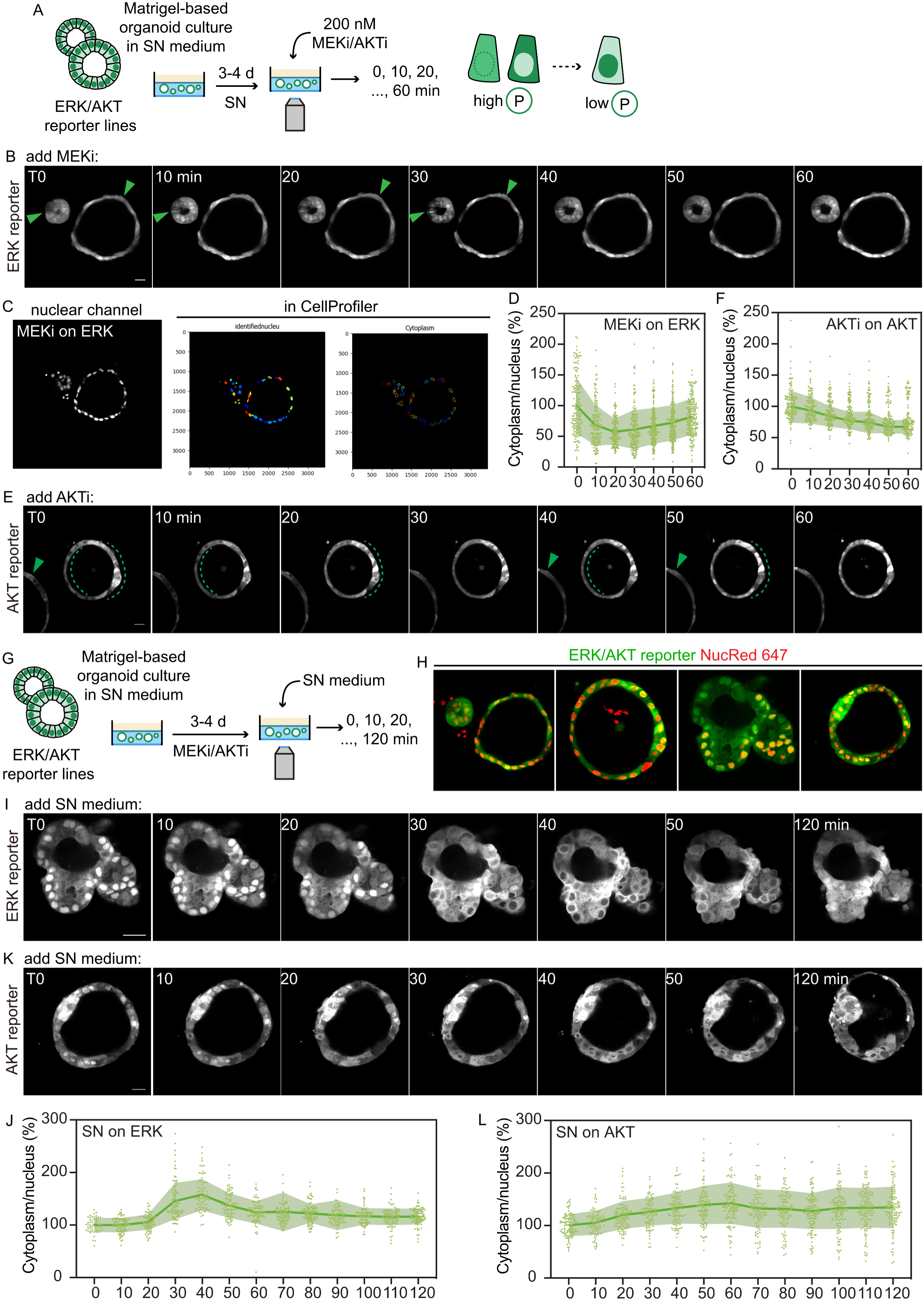
Validation of KTR reporter live-imaging strategy in the organoid system. (Related to Figure 4) (A) Experimental design: ERK or AKT KTR reporter organoids were transferred into a 96- well imaging plate and cultured in SN medium to recover from passaging. MEKi (PD0325901, 200 nM) or AKTi (MK2206, 200 nM) was added to the culture when live-imaging was performed (T0) and the experiment lasted for 60 min. Images of multiple stacks were taken every 10 min during the live-imaging. (B) Representative images showing that the ERK reporter cells responded to MEK inhibition during the live-imaging. Green arrowheads show cells with clear cytoplasm-to-nucleus fluorescence translocation, indicating a reduction in ERK activity. (C) Images showing the quantitation strategy in CellProfiler. The nuclear channel of the live-imaged organoid was segmented (as ‘identifiednucleus’) and the cytoplasm was identified as a grown region (5 pixels) based on the segmented nuclei. Fluorescence intensities for the identified nuclei and cytoplasm of each identified cell (from one image showing one Z plane at one time-point) were recorded. (D) Quantitation of the ERK signal obtained as the cytoplasmic/nuclear intensity for the organoid shown in B. Each point represents one scored cell, dark green line shows the average, light green line shows the 95% confidence intervals. All cytoplasmic/nuclear intensity were normalized to the mean at T0 and shown as a percentage. (E) Representative images showing the AKT reporter cells responding to AKT inhibition during the live-imaging. Green arrowheads and dashed line show cells with clear cytoplasm-to-nucleus fluorescence translocation indicating a reduction in AKT activity. (F) Quantitation of the AKT signal obtained as the cytoplasmic/nuclear intensity for the organoid shown in E. Each point represents one scored cell, dark green line shows the average, light green line shows the 95% confidence intervals. All cytoplasmic/nuclear intensity were normalized to the mean at T0 and are shown as percentage. (G) Experimental design: ERK or AKT reporter organoids in the 96-well imaging plate were treated with MEKi (PD0325901, 200 nM) or AKTi (MK2206, 200 nM) after the cells recovered from passaging. SN medium was added to the culture when live-imaging was performed (T0) and the experiment lasted for 120 min. Images of multiple stacks were taken every 10 min during the live-imaging. (H) Representative images showing the reporter channel (green) and nuclear staining channel (NucRed 647, red) of live-imaged organoids. Representative images showing the ERK reporter cells (I) and AKT reporter cells (K) which received SN medium during the live-imaging. Fluorescence moves from the nucleus to the cytoplasm indicating increased ERK (I) or AKT (K) activity. Quantitation of the ERK signal (J) and AKT signal (L) obtained as the cytoplasmic/nuclear intensity for the organoids shown in H and J, respectively. Each point represents one scored cell, dark green line shows the average, light green line shows the 95% confidence intervals. All cytoplasmic/nuclear intensity were normalized to the mean at T0 and shown as percentage. Scale bars = 20 μm (B, E, I, K).

**Figure S6.**
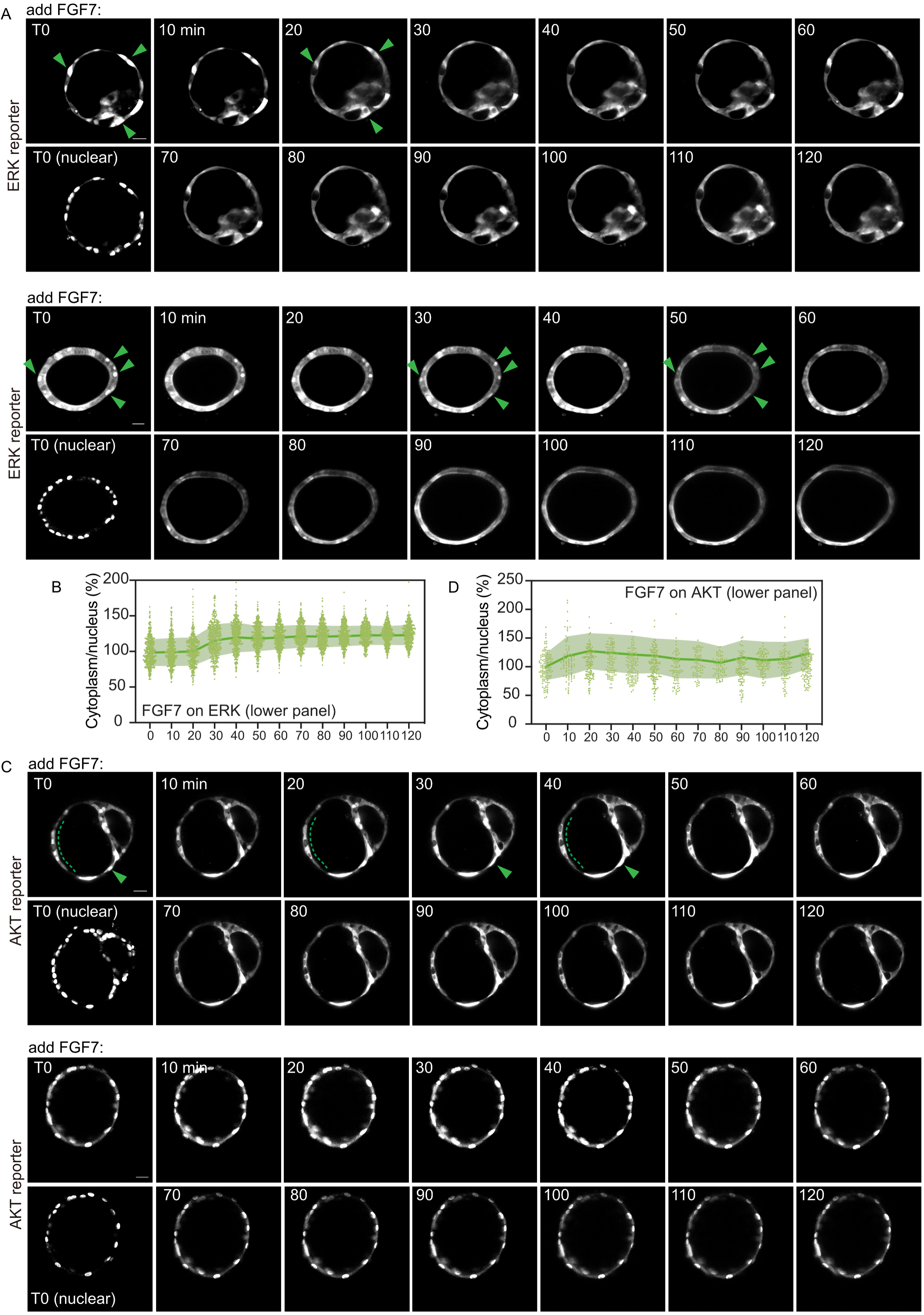
KTR-based live-imaging showing that FGF7 treatment on MEKi- or AKTi-treated organoids restored pERK or pAKT. (Related to Figure 4) Images showing the ERK reporter (A) or AKT reporter (C) cells following FGF7 stimulation. Green arrowheads and dotted lines indicate cells showing evident nucleus-to-cytoplasm fluorescence translocation. (B,D) Quantitation of the ERK signal obtained as the cytoplasmic/nuclear intensity for the organoids shown in A (lower panel) and C (lower panel). Each point represents one scored cell, dark green line shows the average, light green line shows the 95% confidence intervals. All cytoplasmic/nuclear intensity were normalized to the mean at T0 and shown as percentage. Scale bars = 20 μm.

**Figure S7.**
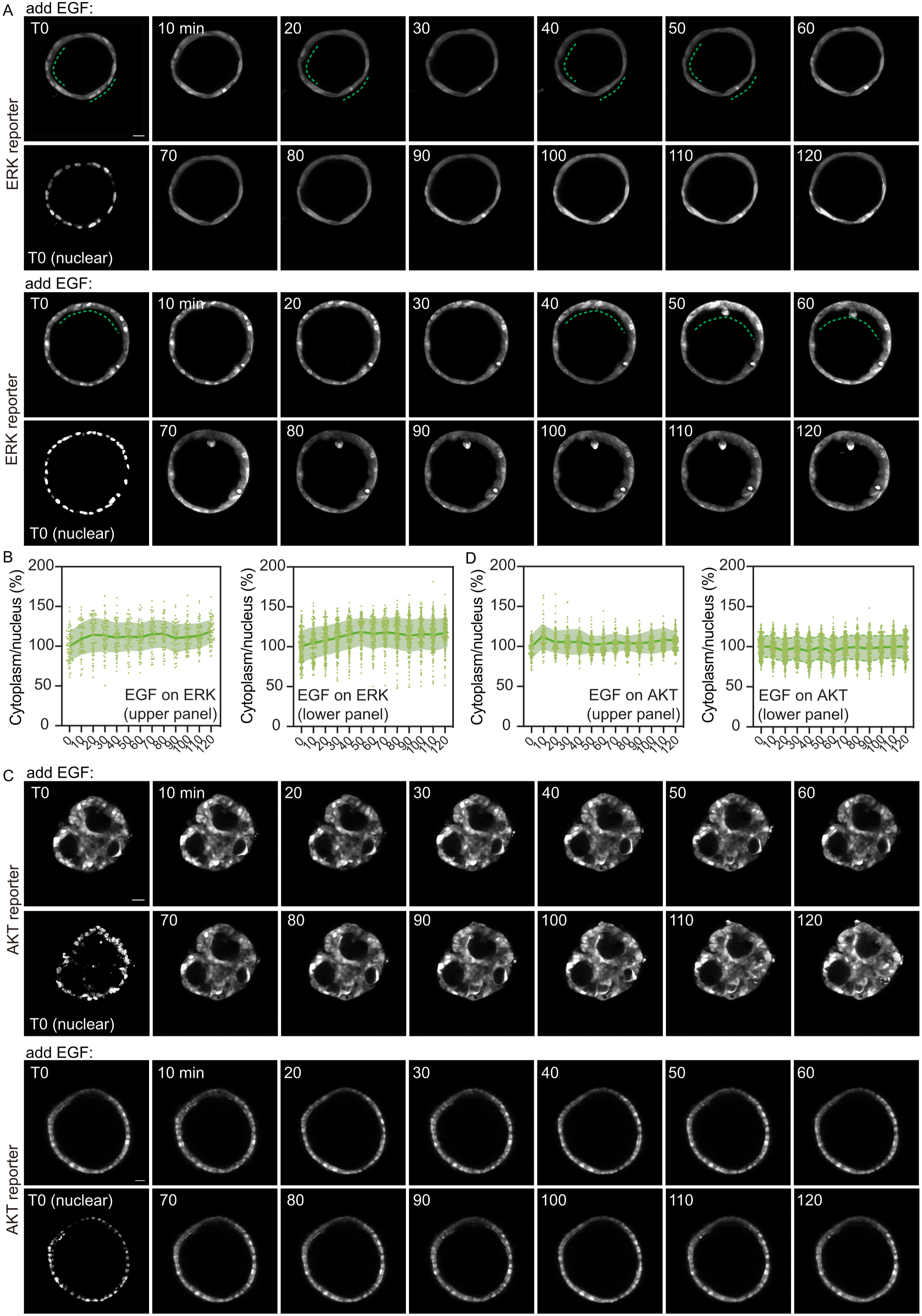
KTR-based live-imaging showing that EGF treatment on MEKi- or AKTi-treated organoids restored pERK, but not pAKT. (Related to Figure 4) Images showing the ERK reporter (A) or AKT reporter (C) cells following EGF stimulation. Green arrowheads and dotted lines indicate cells showing evident nucleus-to-cytoplasm fluorescence translocation. (B,D) Quantitation of the ERK signal obtained as the cytoplasmic/nuclear intensity for the organoids shown in A and C. Each point represents one scored cell, dark green line shows the average, light green line shows the 95% confidence intervals. All cytoplasmic/nuclear intensity were normalized to the mean at T0 and shown as percentage. Scale bars = 20 μm.

**Figure S8.**
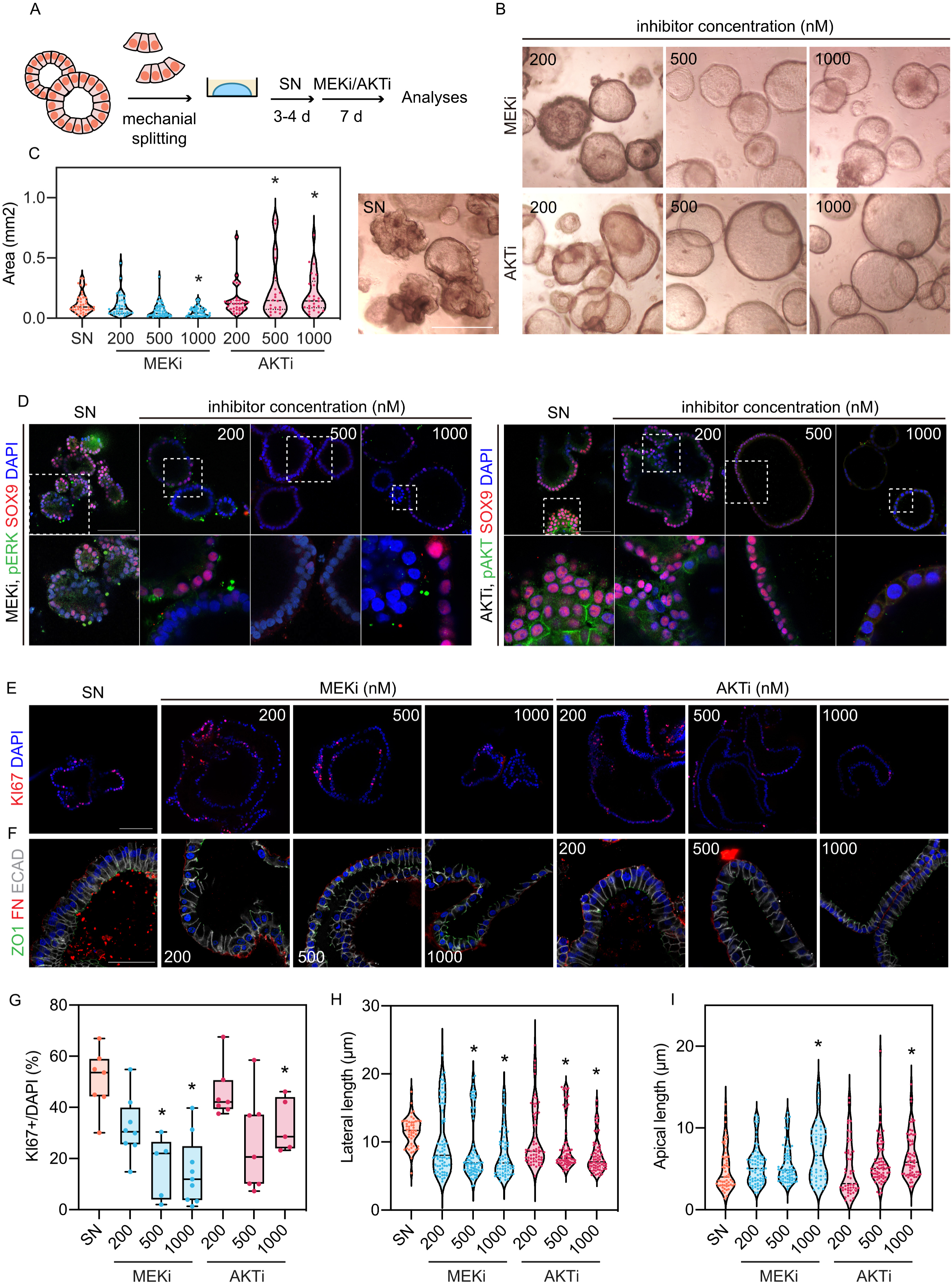
Dose-dependent effects of MEKi and AKTi on the SN organoids. (Related to Figure 5) (A) Experimental design: SN organoids were treated with 200, 500 or 1000 nM MEKi (PD0325901) or AKTi (MK2206) and analysed after 7 days of the inhibition. (B) Representative images showing organoid morphology after 7 days of inhibition. (C) Quantitation of projected area of d7 organoids by in-house Fiji plug-in (Dr. Richard Butler). Mean±s.e.m. are shown. Coloured dots show individual measurements. \**P* < 0.05 (Man- Whitney U test, N = 3 biological replicates). (D) Loss of SOX9 and decreased pERK or pAKT were detected with increasing inhibitor doses. (E) Representative images showing cell proliferation in organoids after 7 days of inhibition. (F) Representative images showing cell shape of organoids after 7 days of inhibition. Quantitation of cell proliferation (G), lateral length (H) and apical length (I) of the organoids related to F. Mean±s.e.m. are shown. Coloured dots show individual measurements. \**P* < 0.05 (Man-Whitney U test, N = 3 biological replicates). Scale bars = 1 mm (B); 100 μm (D, E); 50 μm (F).

**Figure S9.**
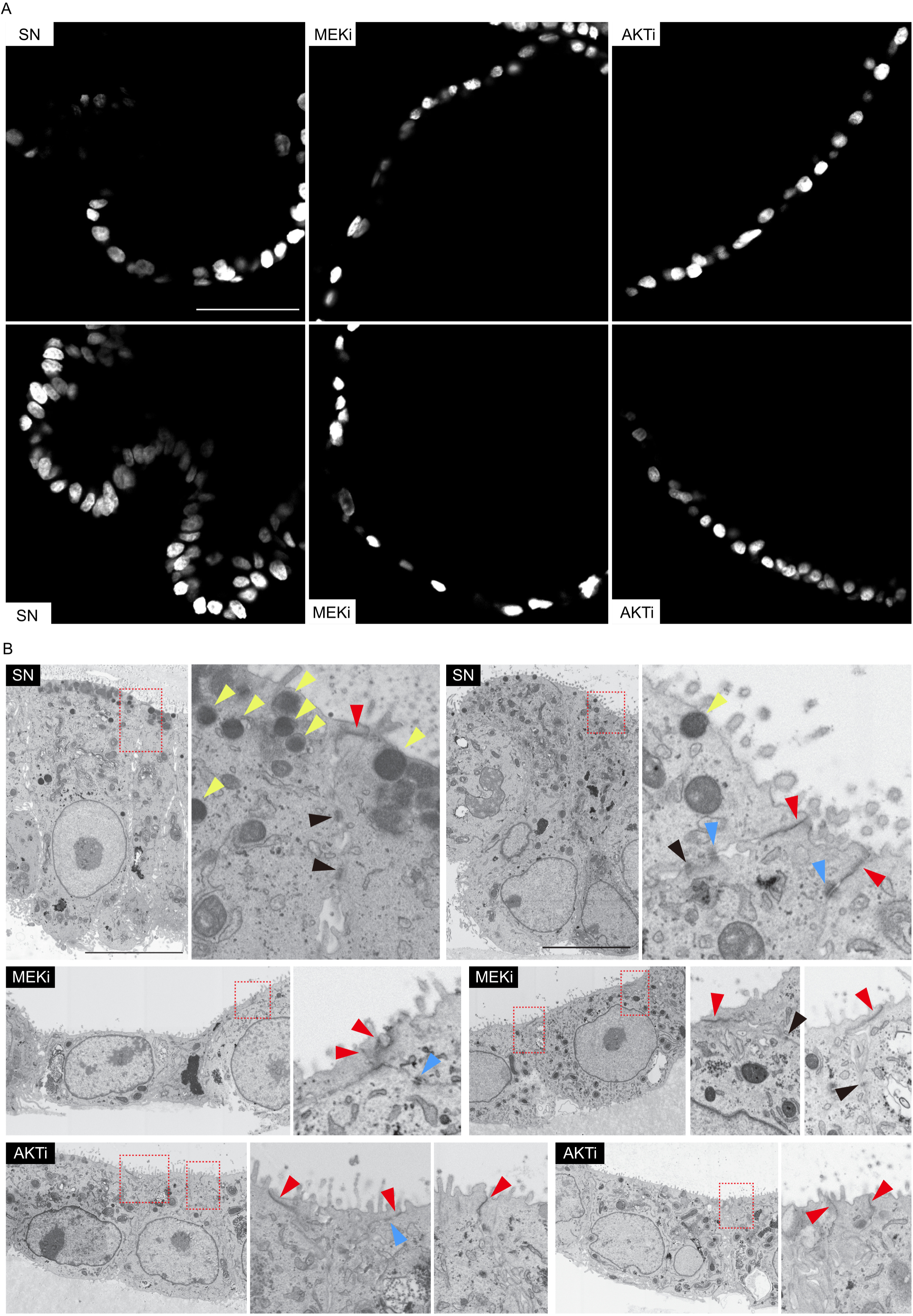
Immunostaining images and SEM images showing cell nuclei, cell structure and junctions in SN, MEKi and AKTi organoids. (Related to Figure 5) (A) Representative images showing cell nuclei of organoids after 7 days of inhibition. Scale bar = 50 μm. Related to Figure 5B and 5H. (B) Representative SEM images showing zonula occluden/adheren (red arrowheads), occludin/claudin (blue arrowheads), adherens junction structure (black arrowheads), and vesicles (yellow arrowheads). Scale bars = 10 μm. Related to Figure 5I.

**Figure S10.**
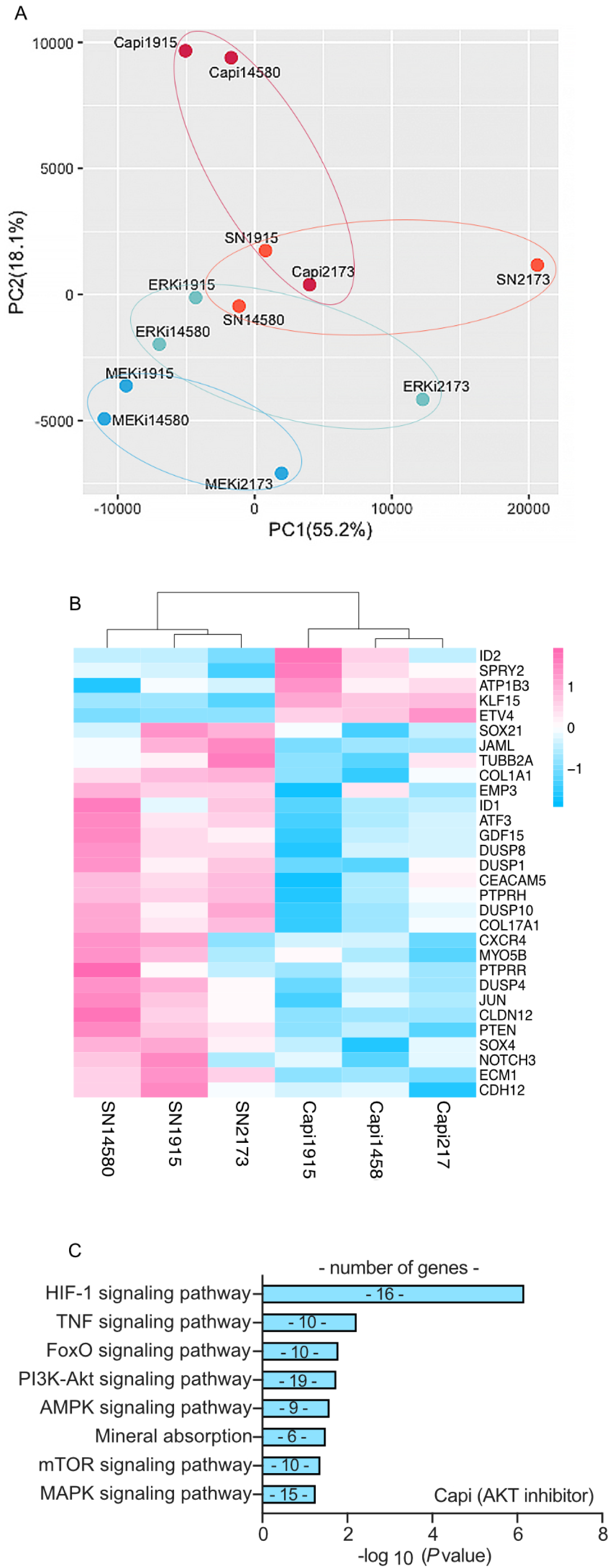
Bulk RNA-seq results showing gene expression variation among organoids treated with different inhibitors. (Related to Figure 6) (A) Principal component analysis of bulk RNA-seq data. (B) Heatmap showing expression level of selected genes significantly altered in AKT-inhibited cells. (C) KEGG pathway analysis (selected terms) of the down-regulated genes in the AKT-inhibited cells; log2FC > 1, adjusted p value < 0.05.

**Figure S11.**
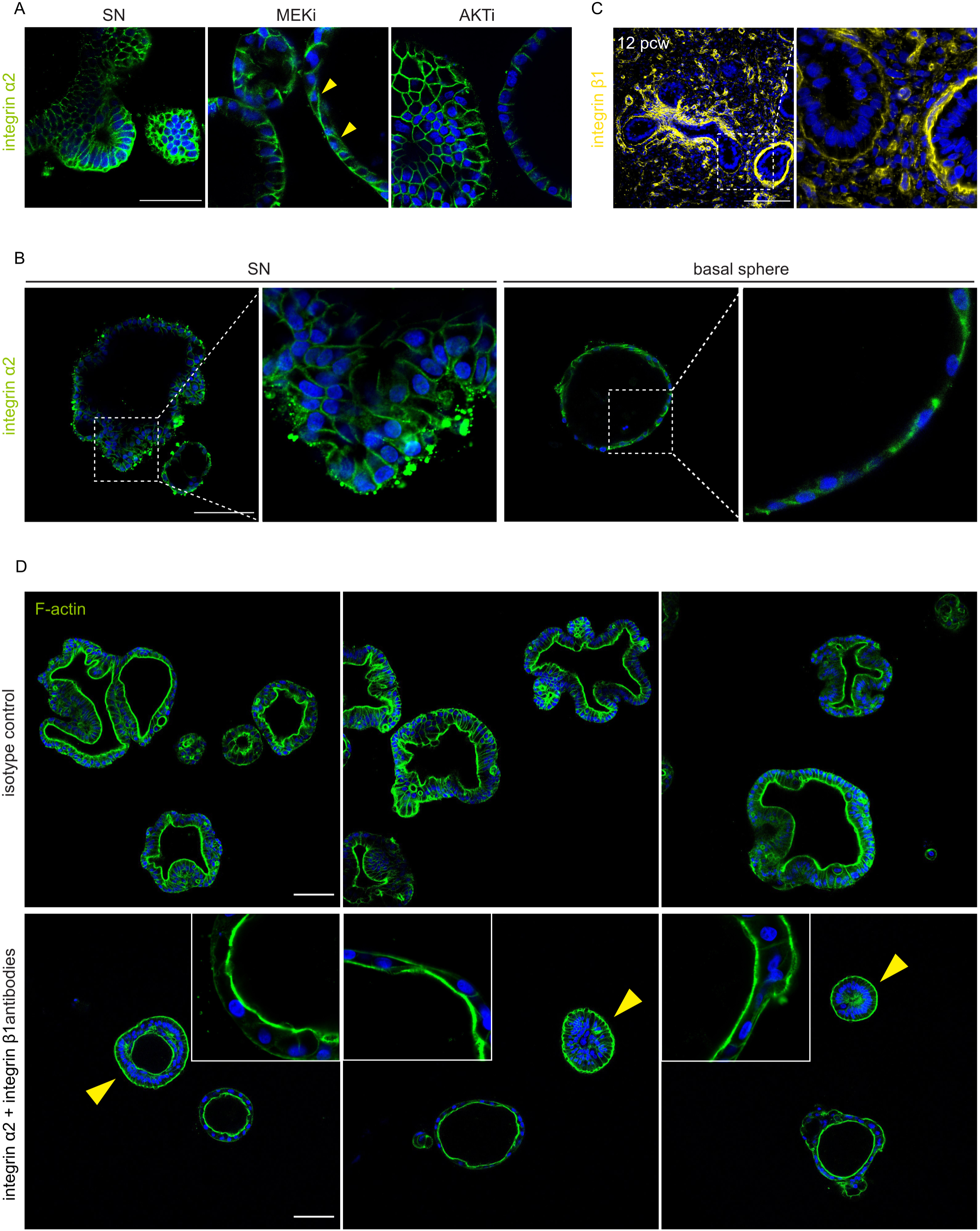
Expression of integrins and organoid morphologies of integrin-blocked cells. (Related to Figure 6) (A) Representative images showing integrin α2 expression pattern in SN organoids and organoids treated with MEKi (PD0325901, 200 nM) or AKTi (MK2206, 200 nM) for 7 days. Yellow arrowheads show altered integrin α2. (B) Representative images showing integrin α2 expression pattern in SN organoids and basal spheres derived from freshly-dissected epithelial tips (as in Fig. 2A) at P1. (C) Representative images showing integrin β1 expression pattern in a 12 pcw human lung. (D) F-actin staining showing organoid morphology, cell shape and cell polarity of organoids treated with isotype control antibody or integrin antibodies. Yellow arrowheads: organoids showing inverted apical-basal polarity. Related to Figure 6F. Scale bars = 100 μm.

**Figure S12.**
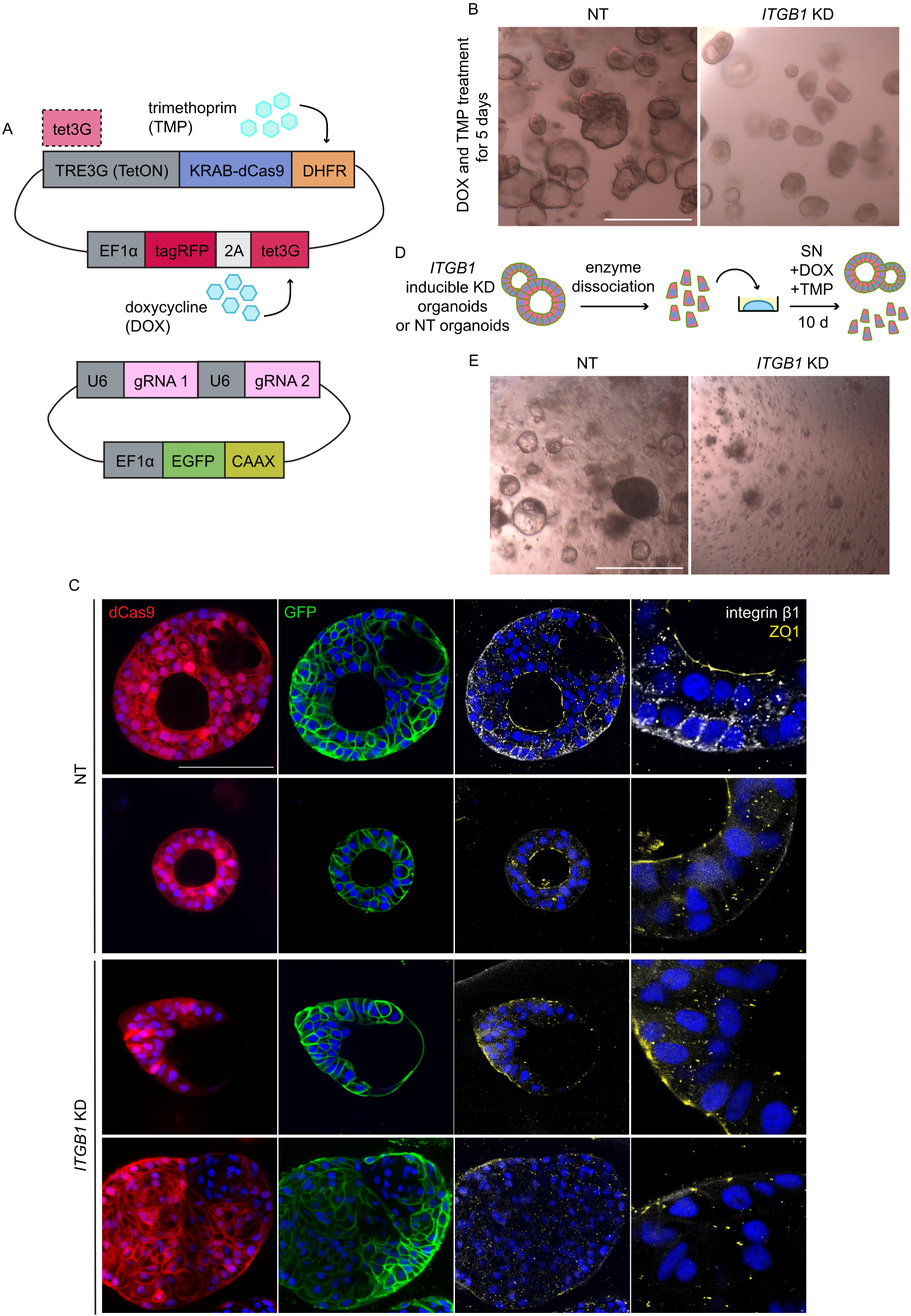
Inducible knockdown of *ITGB1* in the organoids. (Related to Figure 6) (A) Diagram showing the doxycycline inducible KRAB-dCas9 vector and gRNA vector. (B) Representative images showing organoid morphology of *ITGB1*-knockdown (KD) organoid and non-targeting (NT) control after 5 days of 2 μg/ml DOX and 10 μM TMP treatment. (C) Representative images showing organoid morphology, cell shape and polarity of *ITGB1*- KD organoids and NT control organoids after 5 days of DOX and TMP treatment. (D) Experimental design: *ITGB1*-KD organoids or NT control organoids were enzymatically dissociated into single cells and seeded (10,000 per well). 2 μg/ml DOX and 10 μM TMP were added to the single cells for 10 days. (E) Representative images showing organoid morphology of the passaged single cells after 10 days of DOX and TMP treatment. Scale bars = 1 mm (B, E); 100 μm (C).

**Figure S13.**
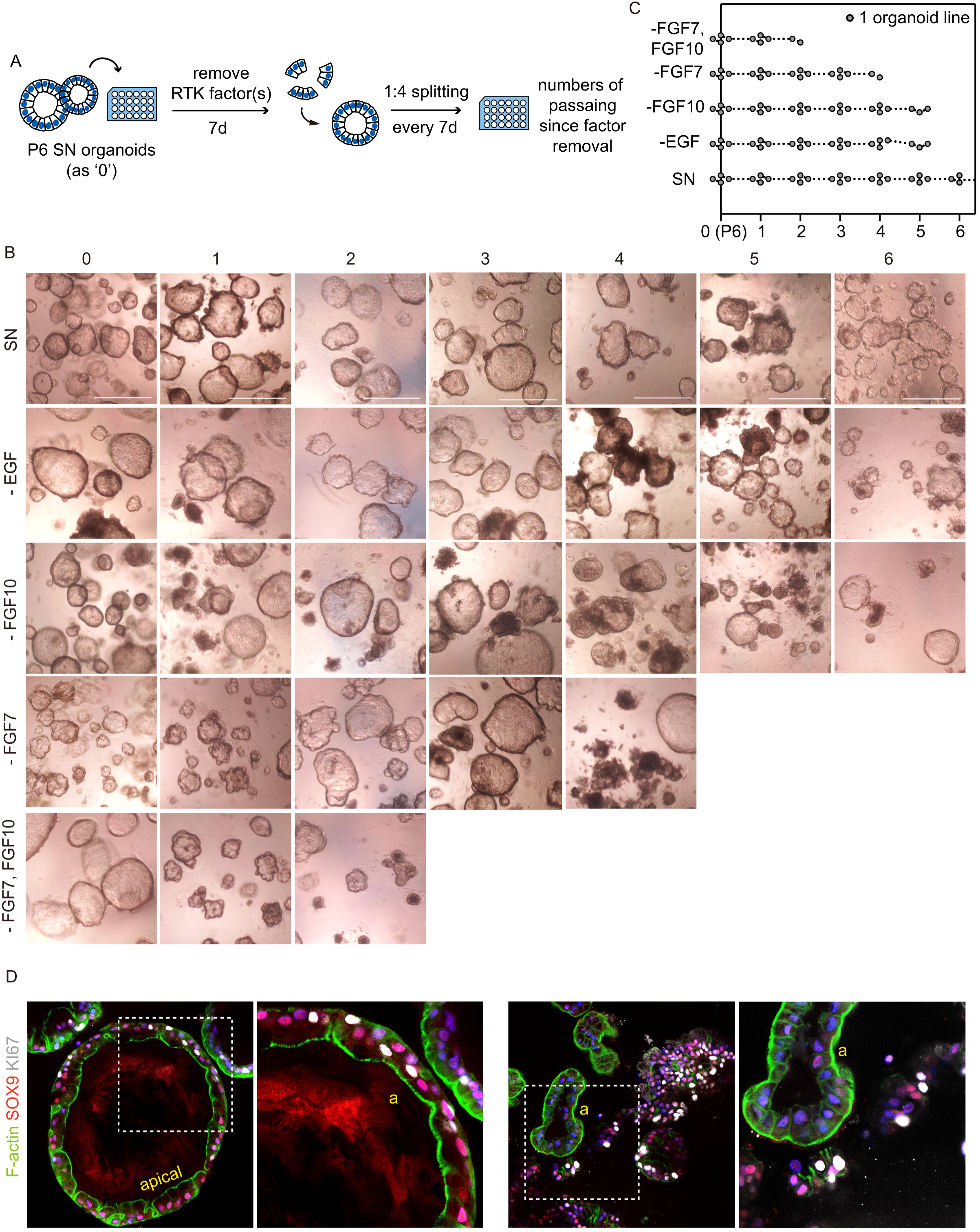
FGF10 is required for the maintenance of SN organoids in vitro. (Related to Figure 7) (A) Experimental design: P6 SN organoids were cultured in media with one or more RTK factors removed from the SN medium. Surviving organoids were passaged every 7d. (B) Representative images showing organoids cultured in different conditions. (C) Survival plot of organoids cultured in each condition. N = 4 organoid lines (8 to 12 pcw) were tested (each point represents one surviving organoid line). (D) Representative images showing organoid morphology and SOX9 expression of organoids in FGF10-removed condition after 5 passages. Scale bars = 1 mm (B); 100 μm (D).

## Notes

### Competing Interest Statement

The authors have declared no competing interest.

